# Functional and evolutionary characterization of potential auxiliary metabolic genes (AMGs) of the global RNA virome

**DOI:** 10.1101/2024.10.16.618626

**Authors:** Yang Zhao, Zhihao Zhang, Pengfei Liu, Meilin Feng, Rong Wen, Yongqin Liu

## Abstract

RNA viruses play an important role in regulating their host and global biogeochemical cycling, and auxiliary metabolic genes (AMGs) are one of the main pathways by which viruses regulate hosts metabolism. However, the AMGs encoded by global RNA viruses are largely unexplored. Here, we annotated RNA viral genomes and identified AMGs from recently published global RNA virome datasets. We found 274 AMGs encoded by 243 RNA viral genomes, which accounts for only 0.008% of all 3,216,257 genomes investigated. These AMGs span 25 functional categories, the most prevalent being translation, energy metabolism, membrane transport, transcription, replication and repair, and signal transduction. Unlike DNA viruses, RNA viruses tend to encode AMGs influencing the genetic and environments information processing of their hosts, with fewer AMGs linked to nutrient cycling, such as carbon and nitrogen metabolism. In addition, RNA viruses encoding these AMGs were taxonomically diverse, spanning five established phyla, the newly proposed phylum “Taraviricota” and fourteen LucaProt supergroups. Host prediction indicated that most of these RNA viruses encoding AMGs infect eukaryotes, while only a few of them, including *Leviviricetes* and *Vidaverviricetes*, infect prokaryotes. Interestingly, the potential host phylum for most of these RNA viruses were different from the likely source phylum of their AMGs, suggesting the possibility that these RNA viruses acquired AMGs from organisms outside their predicted host range. This study provides the first comprehensive analysis of AMGs from global RNA virome, which greatly expands our understanding of their function roles, evolutionary history and virus-host interactions.

## INTRODUCTION

Viruses are the most abundant biological entities on Earth [1], exhibiting immense diversity and playing crucial ecological roles [2–4]. They influence global biogeochemical cycling through close interactions with their hosts. Viruses can reshape microbial communities by lysing host cells [5, 6], thereby altering the abundance and structure of functional microbes. In addition, they modulate ecosystems more directly by altering host metabolism via auxiliary metabolic genes (AMGs) [7]. AMGs are virus-encoded genes acquired from the host and sporadically present in the phage genome to a relatively unknown degree [8, 9]. Viruses randomly sample host genes during infection, and that a subset of these horizontal gene transfer events will be kept in the viral genome if they augment or redirect important metabolic processes that can provide sufficient adaptive advantages to viruses under specific metabolic or nutritional conditions [10, 11].

The discovery of AMGs dates back to the early 2000s, with initial studies focusing on DNA phages infecting marine heterotrophs and cyanobacteria [12, 13]. For decay, research on AMGs has focused on oceanic DNA viruses [9, 11, 14], and the critical role of AMGs in modulating global ocean biogeochemical cycles is well understood [11]. Advancements in metagenomic and metaviromic techniques have expanded the study of AMGs beyond marine environments. DNA viral genomes from a variety of habitats—including humans [10, 15], soils [16–18], wastewater treatment plants [19, 20], and extreme environments [21, 22]—have revealed diverse AMGs with functions ranging from photosynthesis [23], carbon and phosphate metabolism [9, 24], nitrogen and sulfur cycling [14], nucleic acid metabolism [25], and antioxidants [26], and heavy metal detoxification [27].

With the development of metatranscriptomic sequencing, our understanding of the global diversity of RNA viruses has rapidly expanded [28]. Through mining RNA-dependent RNA polymerase (RdRP), the single hallmark protein that is shared by all *Orthornavirae* RNA viruses [29–33], over 10^5^ species of RNA viruses that could be assigned to over 180 RNA viral superclades have been revealed [34]. Prokaryotic RNA viruses, in particular, have surged in number [35–39], with significant implications for Earth’s biogeochemical cycles, given the pivotal role of prokaryotes in driving these processes. Beyond their role in public health, RNA viruses influence ecosystem function by shaping microbial communities, driving microbial evolution, and regulating host metabolism [40–42]. AMGs encoded by RNA viruses are one key mechanism by which they exert these effects [43]. Currently, only five studies have analyzed RNA viral AMGs, all of which focus on specific environments, including one on the cryosphere of the Tibetan Plateau [44, 45], two on marine ecosystems [42, 46] and one on wastewater treatment plants [47], respectively. Despite the availability of global RNA virome datasets covering diverse environments [34, 35, 42], systematic characterization of RNA viral AMGs remains limited. To address this gap and enhance our understanding of the ecological functions, virus-host interactions, and potential evolutionary dynamics of global RNA viruses, we leveraged recently published RNA virus genome datasets— including *Tara* Ocean [42], The RNA Viruses in Metatranscriptomes (RVMT) database [35], and LoucaProt [34]—to perform a functional and phylogenetic analysis of potential RNA viral AMGs.

## RESULTS

### Taxonomy, hosts and environment sources of RNA viruses with AMGs

By screening 3,216,257 RNA viral genomes from global RNA virome datasets including *Tara* Oceans [42], RVMT [35] and LucaProt datasets [34], we identified 274 putative AMGs from 243 RNA virus contigs (vContigs) (Fig. 1; Table S1). Of these RNA viruses, 131 were assigned to five established phyla: *Lenarviricota* (58, 23.9%), *Pisuviricota* (31, 12.8%), *Kitrinoviricota* (30, 12.3%), *Duplornaviricota* (10, 4.1%), and *Negarnaviricota* (1, 0.4%). Additionally, 96 vContigs were classified under the novel “Taraviricota” (1, 0.4%) and 14 distinct LucaProt supergroups (95, 39.1%) (Fig. 2A, B). The LucaProt supergroups primarily consisted of Picorna (37, 15.2%), Tombus-Noda (18, 7.4%), Supergroup022 (8, 3.3%), Yanvirus Supergroup (7, 2.9%), Supergroup044 (7, 2.9%), and Partiti-Picobirna Supergroup (7, 2.9%) (Fig. 2A, B). However, 17 RNA viruses (7.0%) remained unclassified. Among the taxonomically assigned RNA viruses, *Fiersviridae* (35, 14.4%), *Tombusviridae* (14, 5.8%), and *Marnaviridae* (14, 5.8%) were the most represented families (Fig. S1; Table S2).

**Fig. 1.**
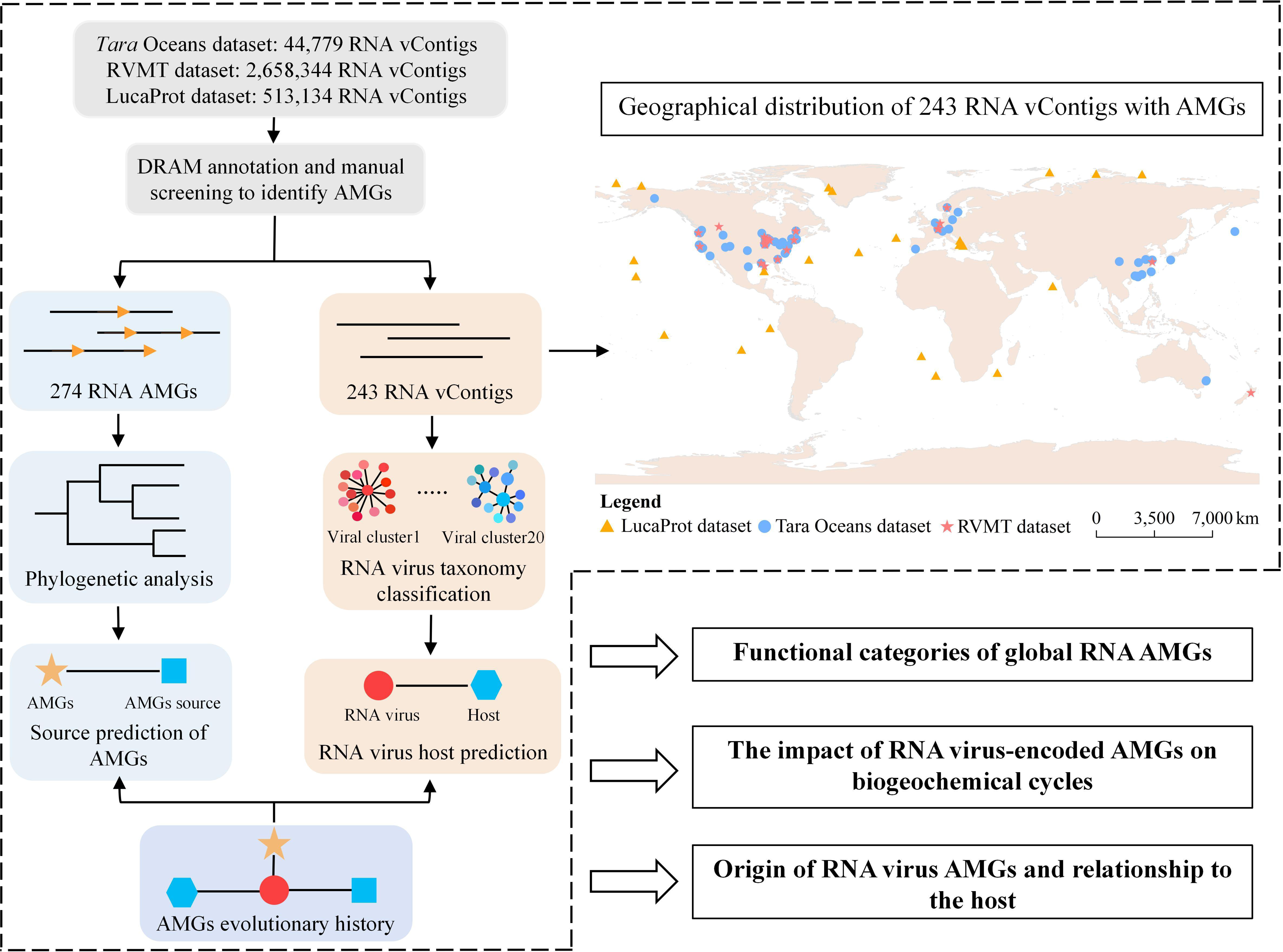
Overview of the global RNA viral AMGs analysis pipeline. The diagram outlines the workflow from global RNA virus genome data collection and processing to the analysis of RNA viral AMGs. 3,216,257 RNA vContigs from 16,408 metatranscriptomes were collected from the *Tara* Oceans, RVMT, and LucaProt datasets to study RNA viral AMGs. Host prediction for RNA vContigs and phylogenetic analysis of AMGs suggested the evolutionary history of selected RNA AMGs and potential virus-host interactions.

**Fig. 2.**
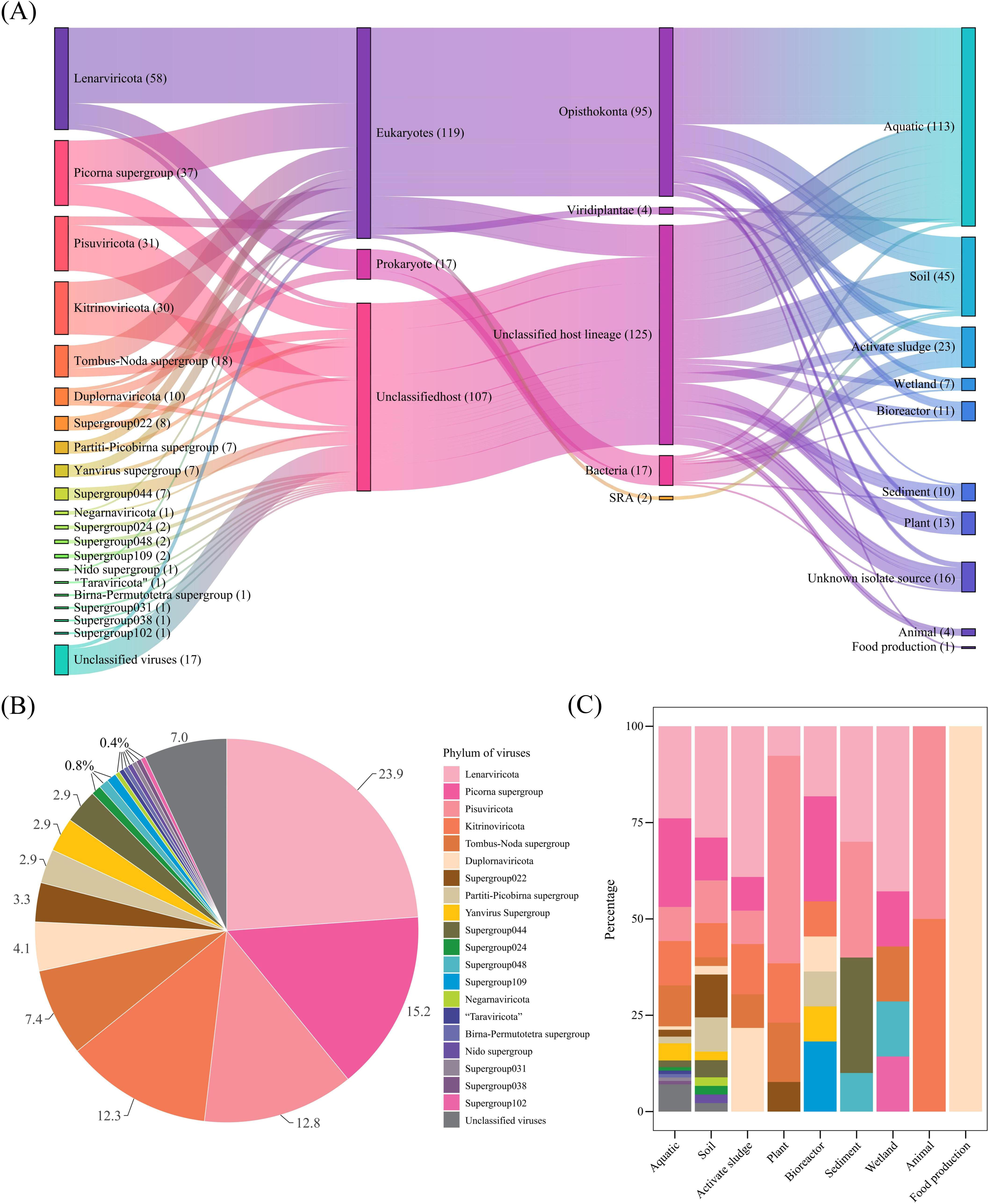
Predicted RNA virus-host interactions. (A) Linkages of RNA vContigs encoding AMGs, their infected hosts, and the environments in which they are detected. The phylum-level taxonomy, predicted hosts and source habitats are shown on the left, the middle two columns and the right columns, respectively. (B) Proportion of RNA vContigs belonging to five established phyla, one “Taraviricota” and 14 LucaProt supergroups. (C) The percentage of each phylum in different habitats and ecosystems. Panels (B) and (C) share the same legend.

Host predictions revealed that RNA viruses with putative AMGs predominantly infect eukaryotes (119, 49.0%), including Opisthokonta (Fungi and Metazoa), the SAR supergroup (Stramenopila, Alveolata, and Rhizaria), and Viridiplantae (green plants). Among these, Opisthokonta emerged as the most frequent eukaryotic host group for these RNA viruses (Fig. 2A; Table S2). Meanwhile, 17 RNA viruses (7.0%) with AMGs belonged to canonical prokaryotic RNA virus groups, including *Leviviricetes* and *Vidaverviricetes* [35]. The hosts of the remaining 107 RNA viruses (44.0%) could not be confidently assigned.

Given that RNA viral genomes from global virome datasets originate from diverse environments [34, 35, 48], we further analyzed the environments from which RNA viruses with AMGs were derived. Aquatic ecosystems (113, 46.5%), soil (45, 18.5%), and activated sludge (23, 9.5%) contained the highest number of vContigs with AMGs (Fig. 2A). *Lenarviricota* had the largest representation in these environments, while the Picorna supergroup was the second most dominant in aquatic and soil habitats, and *Duplornaviricota* ranked second in activated sludge (Fig. 2C). Notably, 10 out of the 17 RNA viruses infecting prokaryotes were identified from activated sludge, likely due to the abundance of bacteria in this environment. In addition, a considerable number of vContigs from plants (13), bioreactors (11), sediment (10), and wetlands (7) were found to encode AMGs. *Lenarviricota* was most abundant in sediments and wetlands, while *Pisuviricota* and the Picorna supergroup were predominant in plants and bioreactors, respectively (Fig. 2C). Although *Kitrinoviricota* was not a dominant phylum, it was widely distributed across various environments, including aquatic ecosystems, soil, activated sludge, plants, bioreactors, and animals.

### Overview of the functions of RNA viral AMGs

Functional annotation revealed that the 274 RNA AMGs were involved in 58 biological pathways (Fig. S2), categorized into metabolic (69, 25.2%), genetic (135, 49.3%), and environmental (35, 12.8%) information processing, as well as cellular processes (14, 5.1%) and organismal systems (3, 1.1%) (Fig. 3). These pathways spanned 25 functional categories, with the top six being: translation (81, 29.6%), energy metabolism (24, 8.8%), membrane transport (23, 8.4%), transcription (19, 6.9%), replication and repair (14, 5.1%), and signal transduction (12, 4.4%) (Fig. 3B, C). The most abundant category was ribosomal proteins (RPs), which are critical for ribosome assembly and protein synthesis. Additionally, RNA AMGs were enriched in genes encoding proteins such as zinc D-Ala-D-Ala carboxypeptidase VanY (7), ABC-2 type transport system permease protein ABC-2.P (5), ATP-binding cassette subfamily C member ABCC1 (5), flagellin FliC (4), molecular chaperone HtpG (4), and cellular nucleic acid-binding protein CNBP (4) (Fig. 3C).

**Fig. 3.**
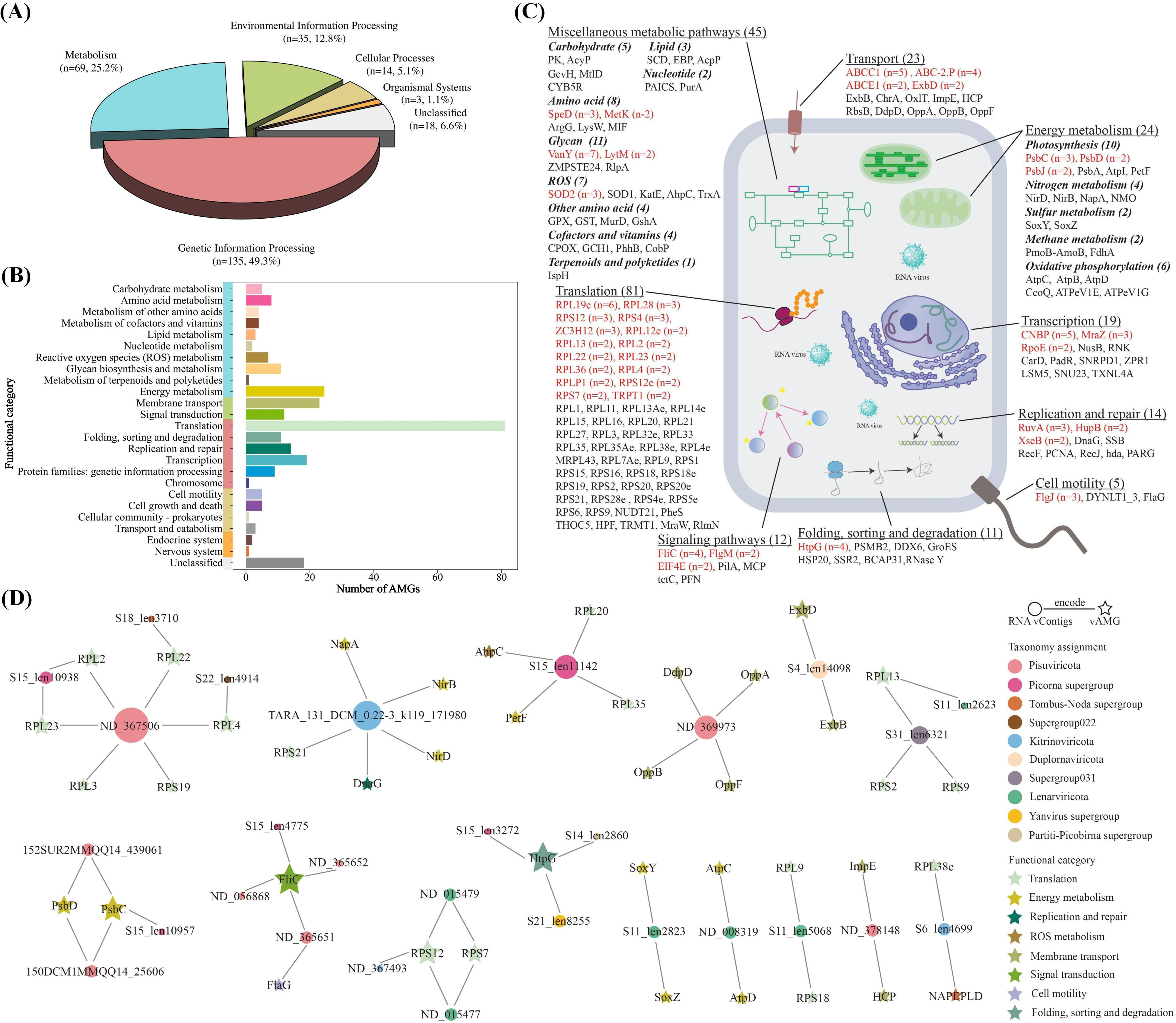
Auxiliary metabolic genes (AMGs) detected in the global RNA virome dataset. (A) The percentage of viral AMGs involved in metabolism, environmental information processing, genetic information processing, cellular processes, and organismal systems. (B) Distribution of 274 RNA viral AMGs in 25 function categories. (C) Number of AMGs associated with main metabolic function categories. (D) Network diagram of 18 RNA vContigs encoding multiple AMGs. Each node represents an vContigs or an AMG, and the color of the node represents the taxonomy of vContigs or the pathway that the AMG is involved in. Each line indicates the vContig encoding the connected AMG.

When comparing the distribution of AMGs across different environments, it was evident that the functional composition varied. RNA viruses from aquatic and soil environments showed enrichment in AMGs related to genetic information processing (translation and transcription) and metabolism, particularly energy metabolism (Fig. S3A, B; Fig. S4). In contrast, RNA viruses from animals, plants, and bioreactors had a higher proportion of AMGs associated with transport and translation (Fig. S4). Additionally, a significant portion of AMGs identified in activated sludge (12, 48.0%), sediments (4, 40.0%), and wetlands (4, 50.0%) were linked to metabolism, where they were enriched in functional categories related to glycan biosynthesis and metabolism, amino acid metabolism and energy metabolism, respectively. (Fig. S3F-I; Fig. S4).

Interestingly, we found that 18 RNA virus contigs encoded more than one AMG. Notable examples include the ribosomal proteins encoded by ND_367506 (*Pisuviricota*) and S15_len10938 (Picorna supergroup), which are functionally connected (Fig. 3D). Furthermore, a vContig from *Kitrinoviricota* (TARA_131_DCM_0.22-3_k119_171980) encoded enzymes involved in nitrogen metabolism (NapA, NirB, and NirD), alongside the ribosomal protein RPS21 and the DNA priming enzyme DnaG, suggesting regulation of nitrogen metabolism, protein synthesis, and DNA replication. Other examples include *Pisuviricota* vContigs 150DCM1MMQQ14_25606 and 152SUR2MMQQ14_439061, which encoded photosynthesis-related proteins PsbC and PsbD; *Lenarviricota* vContig S11_len2823, which encoded sulfur metabolism proteins SoxY and SoxZ; and *Lenarviricota* vContig ND_008319, which encoded oxidative phosphorylation proteins AtpC and AtpD, all of which are central to energy metabolism.

### RNA viral AMGs modulate biogeochemical cycles

To further investigate how RNA viruses may impact biogeochemical cycles, 13 representative AMGs were selected for detailed analysis. These genes are associated with central metabolism (*pk*, *acpP speD*, and *metK*), nitrogen metabolism (*napA, nirB* and *nirD*), sulfur metabolism (*soxY* and *soxZ*), and photosynthesis systems (*psbA, psbC, psbD* and *psbJ*).

### Central metabolism

The first AMG involved in central metabolism is *pk*, which encodes pyruvate kinase (PK), with one instance identified (Fig. 4A). PK is a key enzyme in glycolysis that catalyzes the conversion of phosphoenolpyruvate (PEP) and ADP into pyruvate and ATP. Virus-encoded PK can enhance viral replication by exploiting the host’s metabolic machinery to generate high levels of ATP. The second gene, *acpP*, identified in one RNA vContig, encodes an acyl carrier protein (AcpP) that shuttles acyl intermediates between enzymes during the type II fatty acid elongation process in prokaryotes (Fig. 4B) [49]. The third gene, *metK*, encodes S-adenosylmethionine synthetase (SAM synthetase), which is involved in synthesizing SAM. SAM is an essential chemical for most living organisms and can be used to synthesize a variety of biomolecules, such as proteins, RNA, and phospholipids. In addition, it regulates several biosynthetic pathways such as methionine, polyamine and antibiotic biosynthesis [50]. Two instances of *metK* were identified (Fig. 4C). The fourth gene, *speD*, encodes S-adenosylmethionine decarboxylase [51], which is crucial in spermidine biosynthesis. This enzyme catalyzes the decarboxylation of SAM to produce S-adenosylmethioninamine, which can then be used to synthesize spermidine. Three instances of *speD* were identified (Fig. 4C). Protein structure modeling showed >98% confidence in the prediction of these genes as *pk*, *acpP*, *metK*, and *speD*, respectively (Fig. 4; Table S3).

**Fig. 4.**
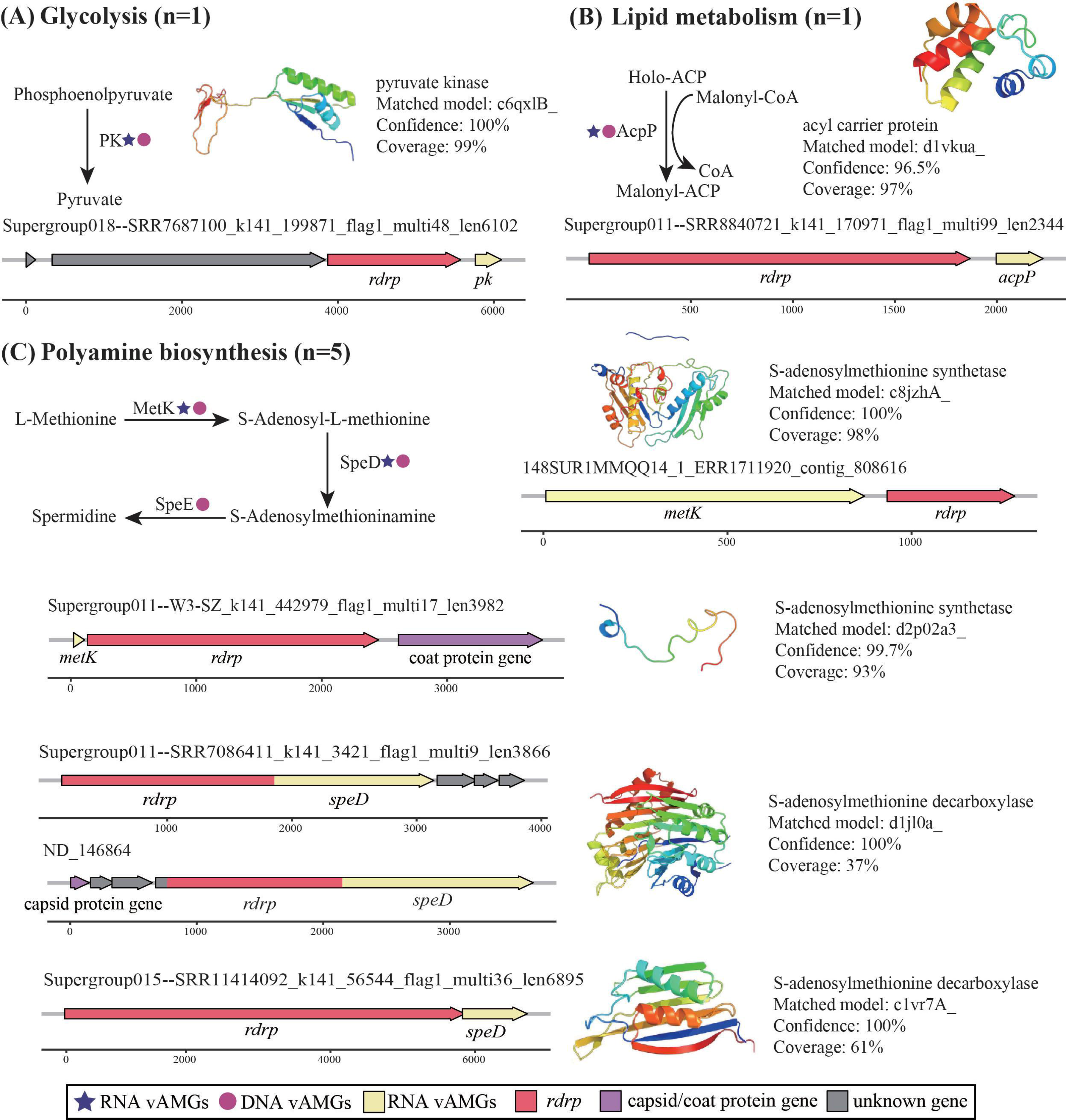
The AMGs encoded by RNA vContigs revealed the involvement of RNA viruses in central metabolism. (A) The metabolic pathways with *pk* gene. The predicted 3D protein structure of PK and the matched reference protein model by Phyre2 are shown next the metabolic pathway. The genome architecture of RNA vContig encoding *pk* is presented at the bottom. (B) Metabolic pathways with *acpP* gene, the reference protein model for the AcpP, and the genome architecture for the RNA vContig encoding *acpP*. (C) Metabolic pathways with *metK* and *speD* genes, reference protein structure model for the MetK and SpeD, and the genome architecture for the RNA vContigs encoding *metK* and *speD* gene. Supergroup011--SRR7086411_k141_3421_flag1_multi9_len3866 and ND_146864 share the same SpeD protein structure model.

### Nitrogen metabolism

The AMGs related to nitrogen metabolism were nitrite reductase *napA*, nitrite reductase (NADH) large subunit *nirB*, and nitrite reductase (NADH) small subunit *nirD*, with only one of each gene identified in RNA viruses (Fig. 5A). NapA, NirB and NirD are members of the dissimilatory nitrate reduction pathway, where NapA catalyzes the reduction of nitrate to nitrite, which can be further reduced to ammonia by NirBD and eventually incorporated into glutamate (Fig. 5A). Protein structure modeling supported the identification of NapA and NirD with >99% confidence (Fig. 5A; Table S3). Although NirB was not confirmed by protein modeling, its full amino acid sequence shares 100% identity with the NCBI reference sequence for NirB (Accession No.: WP_105027197.1) (Table S3).

**Fig. 5.**
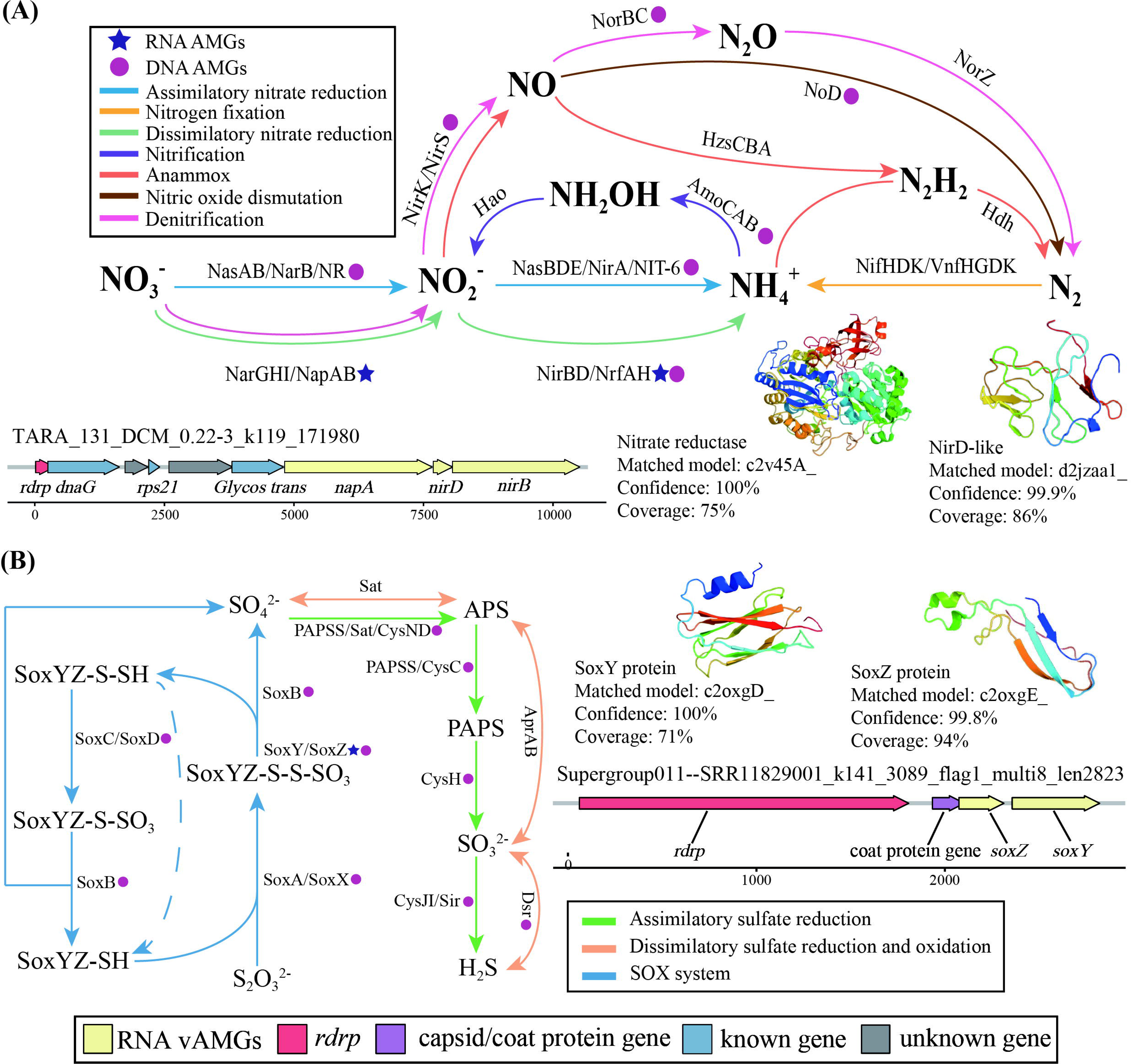
The AMGs encoded by RNA vContigs revealed the involvement of RNA viruses in nitrogen and sulfur metabolism. (A) Potential contribution of AMGs to nitrogen metabolism, genome architecture of RNA vContig encoding nitrogen metabolism genes and reference protein models for NapA and NirD. (B) Potential contribution of AMGs to the sulfur cycling, reference protein models for SoxY and SoxZ, and the genome architecture of RNA vContig encoding sulfur cycling associated genes.

### Sulfur metabolism

The AMGs identified related to sulfur metabolism are the sulfur-oxidizing proteins *soxY* and *soxZ*, each detected once (Fig. 5B). SoxY is a sulfur covalent binding protein that combines with SoxZ to form the SoxYZ complex during sulfur oxidation [52]. Sulfide to the carboxy-terminal cysteine residue of SoxY to form SoxY-cysteine persulfide, which is later oxidized by the sulfane dehydrogenase SoxCD to produce SoxY-cysteine-S-sulfate. Finally, SoxY-cysteine-S-sulfate is hydrolyzed by S-sulfosulfanyl-L-cysteine sulfohydrolase SoxB to produce sulfate [53]. Protein structure modeling confirmed SoxY and SoxZ with >99% confidence (Fig. 5B; Table S3).

### Photosynthesis system

The photosynthetic reaction center PSII (photosystems II) complex is a large protein-pigment assembly in the thylakoid membrane that catalyzes the light-dependent oxidation of water to molecular oxygen [54]. The AMGs involved in PSII are photosystem II protein D1 (*psbA*), photosystem II D2 protein (*psbD*), photosystem II CP43 reaction center protein (*psbC*), and photosystem II reaction center protein J (*psbJ*). Among them, heterodimers composed by D1 and D2 proteins encoded by *psbA* and *psbD* genes are the core of PSII and can bind pigments and cofactors required for primary photochemistry [54]. PsbA and PsbD were found in one and two RNA vContigs, respectively (Fig. 6). The *psbC* gene encodes the chlorophyll binding protein CP43 [55], a light-harvesting system for PSII [56], and was found in three RNA vContigs (Fig. 6). PsbJ, encoded by two AMGs, is not necessary for the photochemical activity of PSII, but plays a role in the stable assembly of water-splitting complex of PSII [57]. The protein structure modeling showed that the confidence of these proteins as PsbA, PsbC, PsbD and PsbJ were all >99% (Fig. 6; Table S3).

**Fig. 6.**
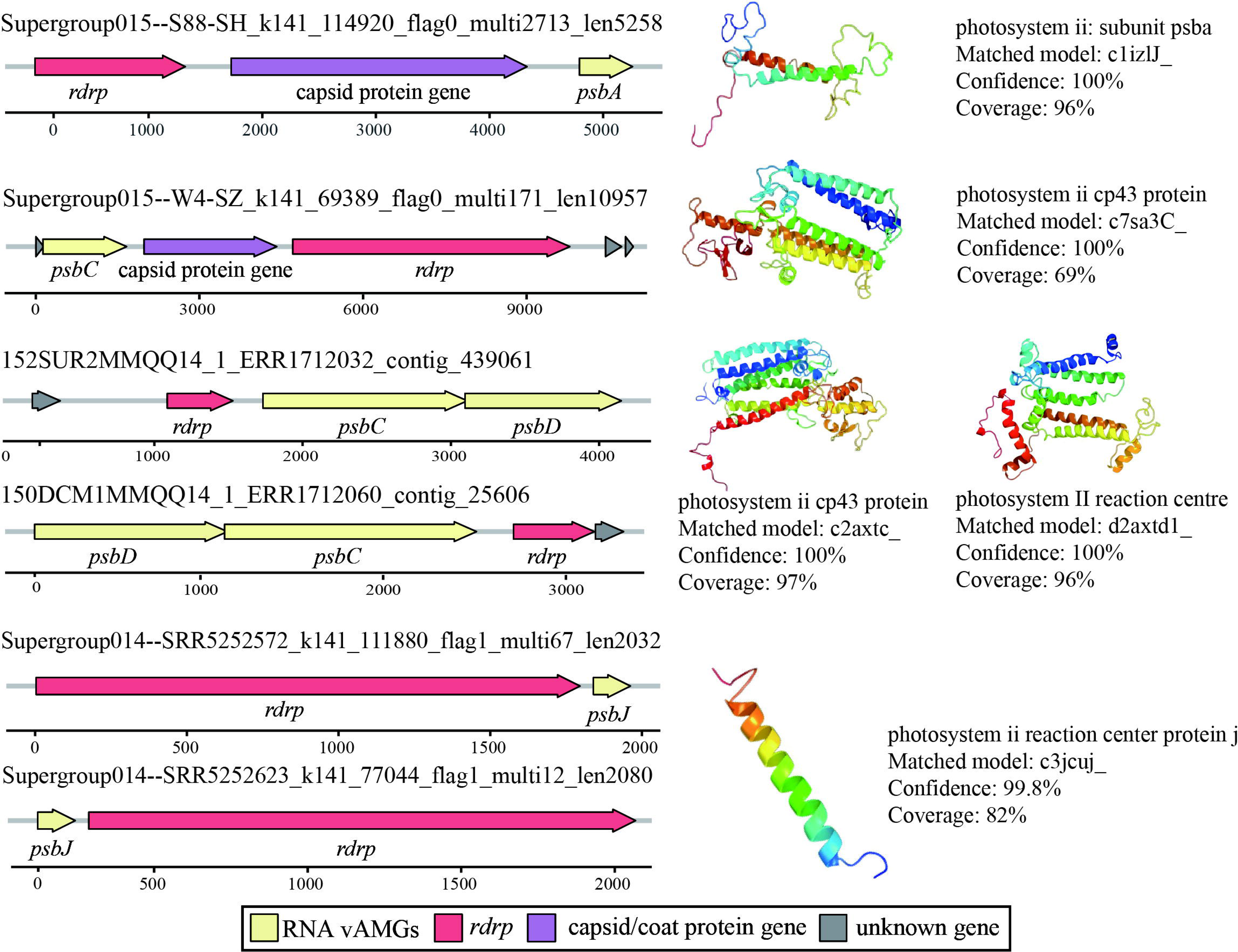
The AMGs encoded by RNA vContigs revealed the involvement of RNA viruses in photosynthesis system. The genome architecture of RNA vContigs encoding *psbA*, *psbC*, *psbD*, and *psbJ* are on the left, and the predictive structure model of the proteins corresponding to these genes is shown on the right. 152SUR2MMQQ14_1_ERR1712032_contig_439061 and 150DCM1MMQQ14_1_ERR1712060_contig_25606 share the same PsbC and PsbD protein structure models. Supergroup014--SRR5252572_k141_111880_flag1_multi67_len2032 and Supergroup014--SRR5252623_k141_77044_flag1_multi12_len2080 share the same PsbJ protein structure models.

### Comparison of the predicted host of RNA viruses and the potential origins of AMGs

To investigate the relationship between the potential sources of AMGs and the hosts of RNA viruses encoding them, we constructed a phylogenetic tree for 30 different types of RNA AMGs (64 sequences from 53 RNA vContigs) associated with biogeochemical cycling, protein synthesis, and motility. Phylogenetic analysis revealed that 38 AMGs encoded by 30 RNA vContigs originated from prokaryotes, while the predicted hosts for 12 vContigs encoding 16 AMG sequences were eukaryotes. Notably, AMGs encoding for PK, AcpP, NapA, NirB, NirD, SoxY, SoxZ, RPL23 (S15_len10938_2), RPL28 (S11_len3419_1), and RPS4 (S11_len4280_5, S11_len4558_1, and S18_len3985_1) are likely derived from Pseudomonadota, although host predictions suggest these RNA viruses infect eukaryotes (Fig. S5; Fig. S6A-J). Additionally, vContigs encoding FliC (S15_len4775_1), FlgM (S11_len4358_1), RPL2 (S15_len10938_1), and RPL4 (S22_len4914_1) infect eukaryotes, yet their AMGs originated from prokaryotic groups such as Thermotogota, Acidobacteriota, Myxococcota, and Verrucomicrobiota, respectively (Fig. S5; Fig. S6K-N). Of the remaining 18 RNA vContigs, three (ND_016429, ND_015477, and ND_015479) were predicted to infect unknown prokaryotes, and 15 infected hosts of unknown identities (Fig. S5; Fig. S6O-P).

Phylogenetic analysis also indicated that 26 AMGs from 24 RNA vContigs were of eukaryotic origin, with one vContig (ND_146864) encoding SpeD predicted to infect prokaryotes. Among the remaining 23 RNA vContigs, 10 were predicted to infect specific eukaryotic phyla, 3 infected eukaryotes of unknown phyla, and 10 infected hosts with unknown identities. Of the 10 RNA vContigs with a predicted host phylum, 7 infected hosts from phyla distinct from the AMG source phylum. For instance, four RNA vContigs encoding MetK (S11_len3982_1), SpeD (S11_len3866_1), RPL13 (S11_len2623_2), and RPL28 (S15_len2589_2) likely originated from Bacillariophyta, yet host predictions suggest they infect Mollusca, Echinodermata, Chordata, and Arthropoda, respectively (Fig. S5; Fig. S6I; Fig. S7A-C). RNA viruses encoding RPL12e (S21_len3745_1) and RPS12e (S15_len3563_1, S15_len2957_2) were predicted to infect Mollusca and Arthropoda, respectively, although their AMGs likely originated from Euglenozoa and Ciliophora (Fig. S5; Fig. S7D-E). Conversely, three RNA vContigs encoding PsbJ (S14_len2032_2, S14_len2080_1) and RPLP1 (S06_len2050_1) were predicted to infect Streptophyta and Ascomycota, respectively, consistent with the phylum of their closest reference sequences of these AMGs in the phylogenetic tree (Fig. S5; Fig. S7F, J). Therefore, only 10% of these investigated RNA vContigs had host phylum that were consistent with the source phylum of their encoded AMGs.

## DISCUSSION

Recent studies have revealed the extensive diversity of RNA viruses across various environments globally [34, 35, 42, 47]. In this study, we provide the first systematic analysis of RNA viral auxiliary metabolic genes (AMGs) on a global scale. Although RNA viruses encode fewer and less diverse AMGs compared to DNA viruses, our results show that RNA viral AMGs still cover a wide range of functional categories, spanning over 25 different functions (Fig. 3). This disparity between RNA and DNA viruses in AMG diversity likely stems from differences in how these genes are acquired. DNA viruses are thought to acquire AMGs directly from the host genome via recombination during infection [10], while RNA viruses likely gain AMGs through RNA-dependent RNA polymerase (RdRp) dissociation events between viral RNA and host mRNA [58]. Since only mRNA transcripts can be transferred and maintained in RNA viral genomes, this limits the types of AMGs RNA viruses can acquire. Additionally, DNA viruses tend to have larger, more flexible genomes and broader host ranges, making gene acquisition easier for them.

DNA viruses are known to encode numerous AMGs involved in key processes such as energy metabolism and biogeochemical cycling of elements, including carbon, nitrogen, phosphorus, and sulfur metabolism [11, 17]. In contrast, RNA viruses appear to have a more limited role in these processes, with only a few AMGs involved in nitrogen and sulfur metabolism, and none related to phosphorus cycling. This suggests that RNA viruses may have a more restricted role in nutrient cycling compared to their DNA counterparts. The host range differences between RNA and DNA viruses might also contribute to this difference, as RNA viruses predominantly infect eukaryotes, while DNA viruses mainly target prokaryotes, including bacteria and archaea.

However, one aspect which RNA viruses excel is in the encoding of ribosomal protein (RP)- related AMGs (Fig. 3C). While cellular organisms encode 102 types of ribosomal proteins, we identified 48 of these in RNA viruses, with the most frequent being RPL19e (6), RPL28 (3), RPS12 (3) and RPS4 (3) (Fig. S8). Ribosomal proteins are crucial for viral protein synthesis, as viruses rely entirely on the host’s translational machinery [59]. After infection, viruses can selectively inhibit the synthesis of host protein by usurping endogenous translation pathways and increasing the biosynthesis of RNA viruses proteins [60, 61]. AMGs are maintained in the virus genome due to their capacity to modify key rate-limiting steps of the host metabolism for the virus cycle, increasing its fitness [42, 62, 63]. The high prevalence of ribosomal proteins in RNA viruses highlights their importance in viral replication, contrasting with the prevalence of energy metabolism and nutrient cycling observed in DNA viruses.

In addition, RNA AMGs can also encode chaperone proteins or peripheral proteins to regulate host environment sensing and stress adaptation. For example, HtpG encoded by RNA AMGs is an ATP-dependent molecular chaperone that is essential for host cell thermotolerance and optimally folds newly synthesized cellular proteins under stress conditions [64]. The ATP-dependent Clp protease (ClpP) can adapt host cells to a variety of stresses by degrading misfolded proteins produced in response to high temperature and stress tolerance (salt, puromycin, ethanol) [65], was found in three RNA vContigs. RNA AMGs involved in flagellar formation can enhance the ability of host cells to adhere, invade, and acquire nutrients by avoiding hostile environments [51]. RNA vContigs can also remodel the host antioxidant network and improve viral replication efficiency by encoding several oxidoreductases (catalase, superoxide dismutase, peroxiredoxin, and thioredoxin) for scavenging reactive oxygen species (ROS) [66].

The close evolutionary relationship between viruses and their hosts suggests that the AMGs encoded by RNA viruses may reflect adaptations to specific environmental conditions. For example, RNA viruses living in the plant phyllosphere have type VI secretion system (T6SS)-related AMGs (HCP and ImpE). RNA viruses carrying these AMGs can help plant hosts to improve their environmental adaptations by inhibiting plant pathogens and pests, and responding to abiotic stresses [67]. Meanwhile, AMGs encoding coproporphyrinogen III oxidase (CpoX), a key enzyme in tetrapyrrole synthesis, were identified in RNA viruses from plant rhizosphere. Since tetrapyrroles are critical compounds for a variety of biological processes (e.g., light-dependent defense and ROS scavenging), RNA viruses with the *cpoX* gene may prevent light-dependent cell death of the plant host and promote its growth [68, 69]. RNA viruses living in mammals enriched with AMGs encoding the oligopeptide transport system, which can mediate the uptake of dipeptides and tripeptides by the animal host to provide it with a nitrogen source [70]. In environments such as soil and activated sludge, RNA viral AMGs encoding antibiotic resistance proteins, like VanY, may reflect adaptation to high concentrations of antibiotics in these habitats [71].

Finally, viruses acquire AMGs from their hosts during infection. Thus, the identified AMGs could also suggest the probable origins of AMGs. For example, RNA viruses encoding photosynthetic proteins likely acquired these genes from Streptophyta (Fig. S5), similar to how cyanophages acquired photosynthesis genes from their cyanobacteria hosts [72].

Interestingly, we found that the potential sources of most of the identified AMGs are different from the predicted hosts of the corresponding RNA viruses encoding them (Fig. S5). The potential hosts of these RNA viruses are assigned according to their taxonomic classification. However, the prediction of uncultured RNA viruses is still a challenging task. Recent studies revealed expanding lineages of prokaryotes RNA viruses [44, 45]. For example, the subsets of picobirnaviruses and partitiviruses known as eukaryote RNA viruses, were most likely to be able to infect prokaryotes [35]. Therefore, these eukaryotes RNA viruses encoding prokaryotes originated AMGs might represent novel lineages of prokaryote RNA viruses. It could also be that these RNA viruses and bacteria infect the same eukaryote hosts, which forms a tripartite association. For example, species diversity of *Bacillus* as endophyte and rhizobacteria of cotton plants was associated with tobacco streak virus (TSV) infection in cotton [73]. During the co-infection, RNA viruses of eukaryotes obtained bacteria encoded genes. In addition, it could be that the RNA viruses AMGs is acquired by their hosts through HGT from a third organism. Lastly, we acknowledge that some RNA viral contigs may be the result of misassembly. The RNA viral contigs in this study were assembled from short reads using *de novo* assembly, and while the risk of chimeric assemblies is low, it cannot be entirely ruled out [74]. Future studies using long-read sequencing or attempts to isolate these RNA viruses will help resolve these uncertainties and further elucidate the role of RNA viral AMGs in host-virus interactions.

## Limitations

During the preparation of this work, studies on RNA viruses of the cryosphere [44, 45], wastewater treatment plants (WWTP) [47] are published as preprints, which are not included in the current paper. Compared with these studies, similar patterns were observed for AMGs encoded by RNA viruses from these environments. Specifically, Sun et al. analyzed RNA AMGs from the coastal region of Qingdao and identified 36 AMGs, which were mainly associated with signaling pathways, membrane transport, and transcriptional pathways [46]. Yuan et al. found that RNA AMGs in global WWTP are involved in diverse host metabolic processes, mainly including improved translation efficiency, cellular respiration, nitrogen metabolism, and even antibiotic resistance [47]. Moreover, photosynthesis and protein synthesis related AMGs were widely detected in glacier and ocean environments [42, 44]. The widespread distribution of these AMGs emphasizes their importance in regulating the metabolism and virus-host interactions of RNA viruses. Compared to the published datasets of RNA viral AMGs [42, 44–47], 75.1% (145 out of 193 different types of AMGs) of the AMGs identified in this analysis are new, which significantly expands the existing diversity of global RNA viral AMGs. Currently, only a small portion of environments of the global ecosystems are covered by RNA virome studies [75, 76]. Together with our synthesis, we predict that a greater diversity of RNA viral AMGs will be uncovered with more comprehensive sampling of the global RNA virome.

## CONCLUSION

By leveraging global RNA virome datasets, we generated the first comprehensive view of RNA viral auxiliary metabolic genes (AMGs). Our findings revealed several key insights: (i) RNA viruses exhibit remarkably high AMG diversity, spanning 25 distinct functional categories, with the most prevalent functions related to translation, energy metabolism, transport, and transcription; (ii) RNA AMGs predominantly encode proteins involved in the regulation of environmental and genetic information processing, while those associated with nutrient cycling are less common; (iii) RNA viruses carrying AMGs are capable of infecting both eukaryotes and prokaryotes, with *Leviviricetes* and *Vidaverviricetes* being the most common groups infecting prokaryotes; (iv) the presence of RNA AMGs encoding antibiotic resistance, anti-oxidative stress, chaperonins, and periplasmic proteins suggests a role in helping hosts adapt to specific environmental conditions; and (v) RNA viruses may acquire AMGs from organisms outside their predicted host range. Together, these findings greatly advance our understanding of the complex interactions between RNA viruses, their hosts, and the environment.

## METHODS

### Global RNA virome genome data collection

We compiled a global RNA viral contig (vContigs) dataset comprising 3,216,257 RNA vContigs from three major sources: the *Tara* Oceans RNA viral dataset (TO), The RNA Viruses in Metatranscriptomes (RVMT) database, and the LucaProt RNA virome dataset, all previously published [34, 35, 42]. The TO dataset represents the first global marine RNA virus catalog, derived from 771 marine metatranscriptomes (187 prokaryote-enriched and 584 eukaryote-enriched) sampled from 121 oceanic stations during the *Tara* Oceans and *Tara* Oceans Polar Circle expeditions. A total of 44,779 RNA vContigs were identified from these metatranscriptomes. The RVMT dataset encompasses a broader environmental scope, covering aquatic, marine, freshwater, soil, plant, animal, and engineered systems, and includes 5,150 metatranscriptomes, from which 2,658,344 RNA vContigs were identified. Finally, the LucaProt dataset spans 10,487 metatranscriptomes from diverse ecosystems, identifying 513,134 RNA vContigs through a deep-learning-based analysis using the LucaProt algorithm.

### Identification of the auxiliary metabolic genes (AMGs)

Auxiliary metabolic genes (AMGs) encoded by RNA vContigs were identified through a combination of functional annotation and manual curation. Initially, all 3,216,257 RNA vContigs were annotated using the Distilled and Refined Annotation of Metabolism (DRAM v1.4.6) pipeline with default parameters [77]. DRAM is a tool for genome annotation, and KOfam, UniRef-90, PFAM-A, dbCAN, RefSeq viral, VOGDB, and the MEROPS peptidase databases were used for homolog search and genome annotations in the current study. Following DRAM annotation, we manually filtered the results to focus only on genes involved in cell-associated metabolic processes, which were classified as AMGs. Genes encoding conserved viral proteins, such as structural proteins (e.g., viral coat/core and pilin-like proteins) and replication/assembly proteins (e.g., DNA/RNA polymerases and nucleotide-binding proteins), were excluded [78]. Finally, To further confirm the functions of the identified AMGs, we applied the Phyre2 pipeline (http://www.sbg.bio.ic.ac.uk/phyre2) for structural analysis [79], while AMGs with confidence scores below 90% were subjected to additional functional validation by comparison with similar sequences in the NCBI nr database (Release May, 2024). The 274 RNA AMGs obtained were compared with five published repositories of RNA AMGs [34, 42, 44, 45, 47], which we regarded as potentially novel when the gene, enzyme and KO names were different from previously identified.

### Domain and functional annotation of RNA viral genomes with AMGs

The 243 RNA vContigs with AMGs were further subjected for domain annotation by using three methods: Open reading frames (ORFs) based on default code. The nucleotide sequences of 243 vContigs were annotated InterProScan (v5.62-94.0) [80] against PRINTS v.42.0 [81], Phobius v.1.0.1 [82], MobiDBLite v.2.0 [83], and TMHMM v.2.0c [84] database.

i. ORFs based on optimized code. The optimization code was determined by the longest sequence translated by transeq (EMBOSS v.6.5.7.0) and its corresponding code table. For RNA viral genome domain annotation, ORFs were identified by using Prodigal v2.6.3 with the optimized genetic code. Profiles of amino acid sequences were generated using HHsuite v3.3.0 [85] and UniRef30_2020_03 [86] database with one iteration of hhblits under the option “-n 1 -e 0.001”. Then, Pfam 35 was used to identify structural domains in the amino acid sequences and the domains annotated using hits with >95% probability score [48].
ii. Domain-based. Since there is still a challenge to determine the use of the code, we also performed functional domain annotation on the translated RNA vContigs sequences. Protein sequences from contigs translated by transeq for each RNA vConitgs were annotated by hmmsearch [87], which matched these proteins to Hidden Markov Models (HMM) collected from multiple protein mapping databases (CDD v.3.19 [88], Pfam 35 [89], Gene3D v4.3 [90], LysDB [35], ECOD 2017 release [91], with a maximum E-value of 0.001.

The genome architecture of RNA vContigs with AMGs was visualized using gggene (https://github.com/wilkox/gggenes).

### Taxonomy and host prediction of RNA viruses with AMGs

In this study, we utilized RNA virus species and host information from Dominguez-Huerta et al., Zayed et al., Neri et al., and Hou et al., which were primarily predicted based on RNA-dependent RNA polymerase (RdRp) sequence similarity and RNA virus taxonomy [34, 35, 42, 48]. The TO dataset identified 5504 “species”-level viral operational taxonomic units (vOTUs) belonging to 10 phyla, including five established and five new proposed phyla (“Taraviricota”, “Pomiviricota”, “Arctiviricota”, “Paraxenoviricota” and “Wamoviricota”) [48]. These RNA viruses are known to infect various ecologically important organisms, primarily protists and fungi, with fewer infecting bacteria and invertebrate metazoans [42]. In the RVMT dataset, RNA viral diversity expanded from 13,282 to 124,873 distinct clusters at a taxonomic level between species and genus. Additionally, two candidate phyla (p.0001.base-Kitrino and p.0002.base-Kitrino) were proposed alongside the five previously established phyla. This dataset revealed that most RNA viruses are associated with eukaryotic hosts, with only two groups, leviviruses (*Leviviricetes*) and cystoviruses (*Vidaverviricetes*), known to infect bacteria [35]. The LucaProt dataset identified 161,979 putative viral species and 180 RNA viral supergroups, many of which are comparable to known virus phyla and classes defined by the International Committee on Taxonomy of Viruses (ICTV). Of these supergroups, only 21 corresponded to existing ICTV-classified viral phyla/classes, while 60 of the remaining 159 represented highly divergent, previously unrecognized supergroups. Though many of these RNA viruses likely associate with diverse microbial eukaryotes, a significant proportion of the newly discovered viruses are predicted to infect Bacteria and potentially Archaea [34].

### Phylogenetic analysis of RdRp and AMGs

We constructed phylogenetic trees of RdRp amino acid (AA) sequences encoding the four RNA virus phyla with the highest number of AMGs, which were classified into *Lenarviricota*, Picorna supergroup, *Pisuviricota* and *Kitrinoviricota*. Reference sequences were obtained from the TO, RVMT and LucaProt datasets. Briefly, reference RdRps and RDRPs of RNA viruses with AMGs that from the same phylum were pooled together and aligned using MUSCLE5 with default parameters [92]. The comparison sequences are filtered using trimAl V 1.5.0 (-gappyout parameter) [93] and a custom script (fasta_drop.py) to remove sequences with >70% gaps. The final multiple sequence comparison results were used for maximum likelihood tree construction using MEGA software [94, 95]. The tree topology was evaluated using the bootstrap analysis based on 1,000 resampling replicates.

To investigate the probable evolutionary origin of the RNA AMGs, homolog sequences of RNA AMGs were recruited from the NCBI nr database (Release May, 2024), and the top 20 hits with a bit score threshold of 50 were retained as reference sequences for the RNA AMGs. The RNA AMGs and reference sequences were aligned using MUSCLE5, and manually checked and trimmed. The phylogenetic trees were constructed using the methods described above and visualized using the iTOL online server [96].

## Authors’ contributions

PL and YL initiated the study; PL and YL designed the study; YZ, ZZ and PL performed all the data analysis with results interpretations from also MF and RW; YZ and PL led the manuscript writing with the help from ZZ, MF and RW. All authors contributed to the editing of the text and approved the final version.

## Acknowledgment

This work was supported by the National Natural Science Foundation of China General Program (42171144), the National Natural Science Foundation of China for Excellent Young Scientists Fund Program (42222105), the Second Tibetan Plateau Scientific Expedition and Research Program (STEP) (2021QZKK0100), and the Global Ocean Negative Carbon Emissions (Global ONCE) Program.

