## Supplementary Figures for "Functional and evolutionary characterization of potential auxiliary metabolic genes (AMGs) of the global RNA virome"

Yang Zhao et al.

**This PDF file includes:**

Supplementary Figs. 1 to 8

**Other Supplementary Materials for this manuscript include the following:**

Supplementary Table S1 to S4 in one Excel file.

### Supplementary figures

(A)

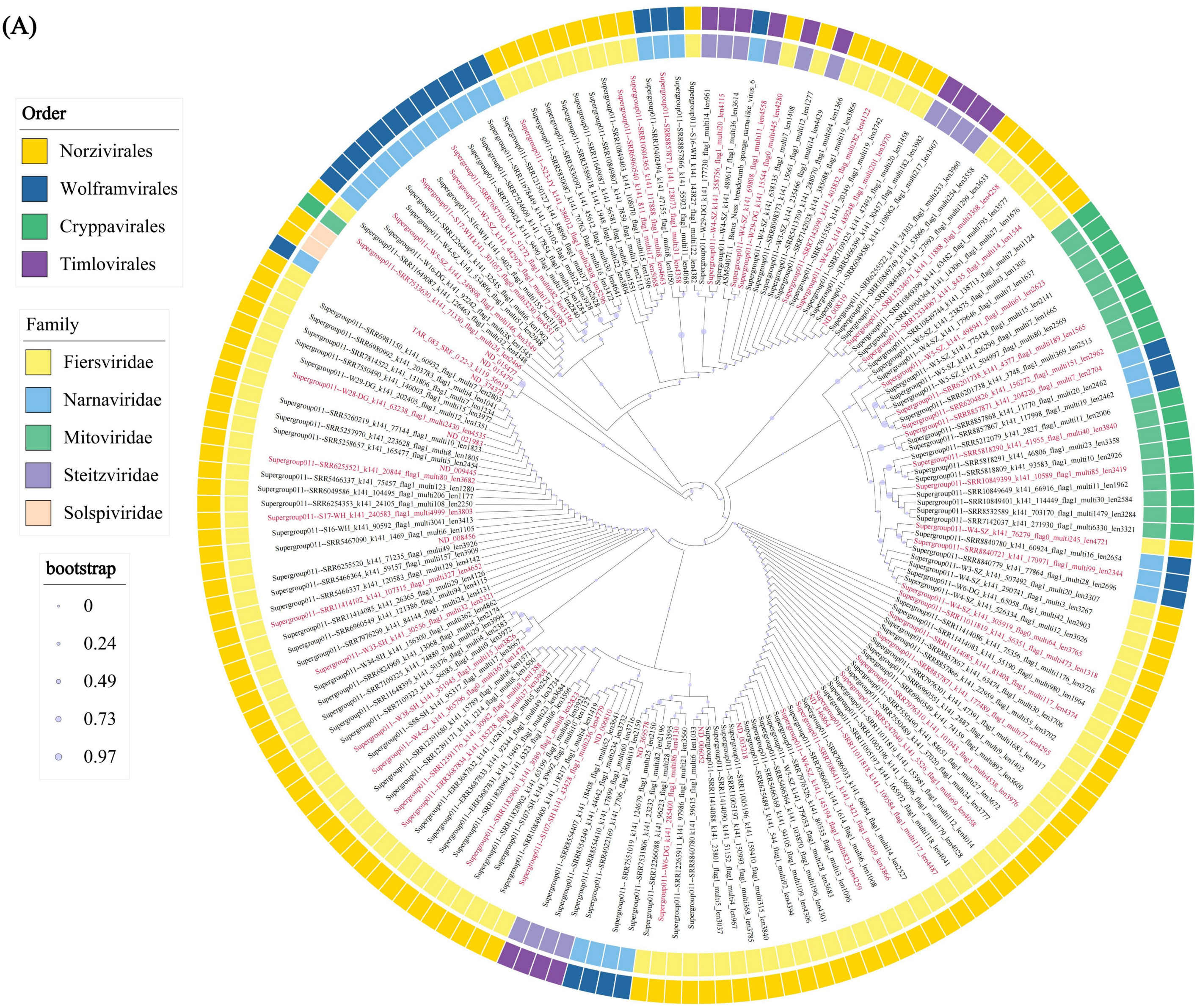


**Fig. S1 | Phylogenetic tree of RdRps for four phylum. (A)** Phylogenetic tree of RdRps for the Lenarviricota phylum. RNA viral RdRPs with AMGs are highlighted in red, while reference sequences are highlighted in black. The outer ring shows the order and family of RdRps, respectively.

**(B)**

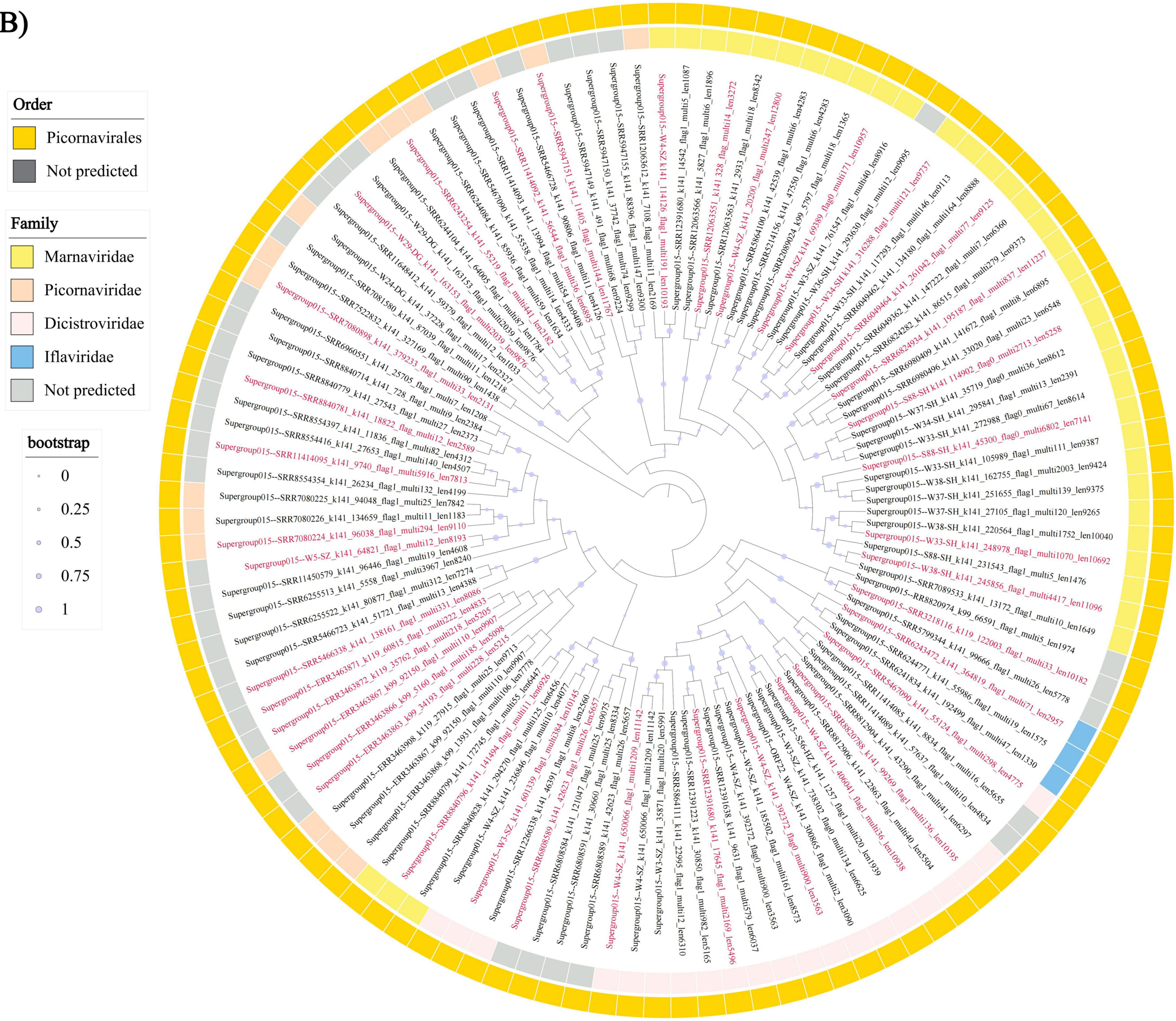


**Fig. S1 | Continue. (B)** Phylogenetic tree of RdRps for the picorna supergroup phylum. RNA viral RdRPs with AMGs are highlighted in red, while reference sequences are highlighted in black. The outer ring shows the order and family of RdRps, respectively.

(C)

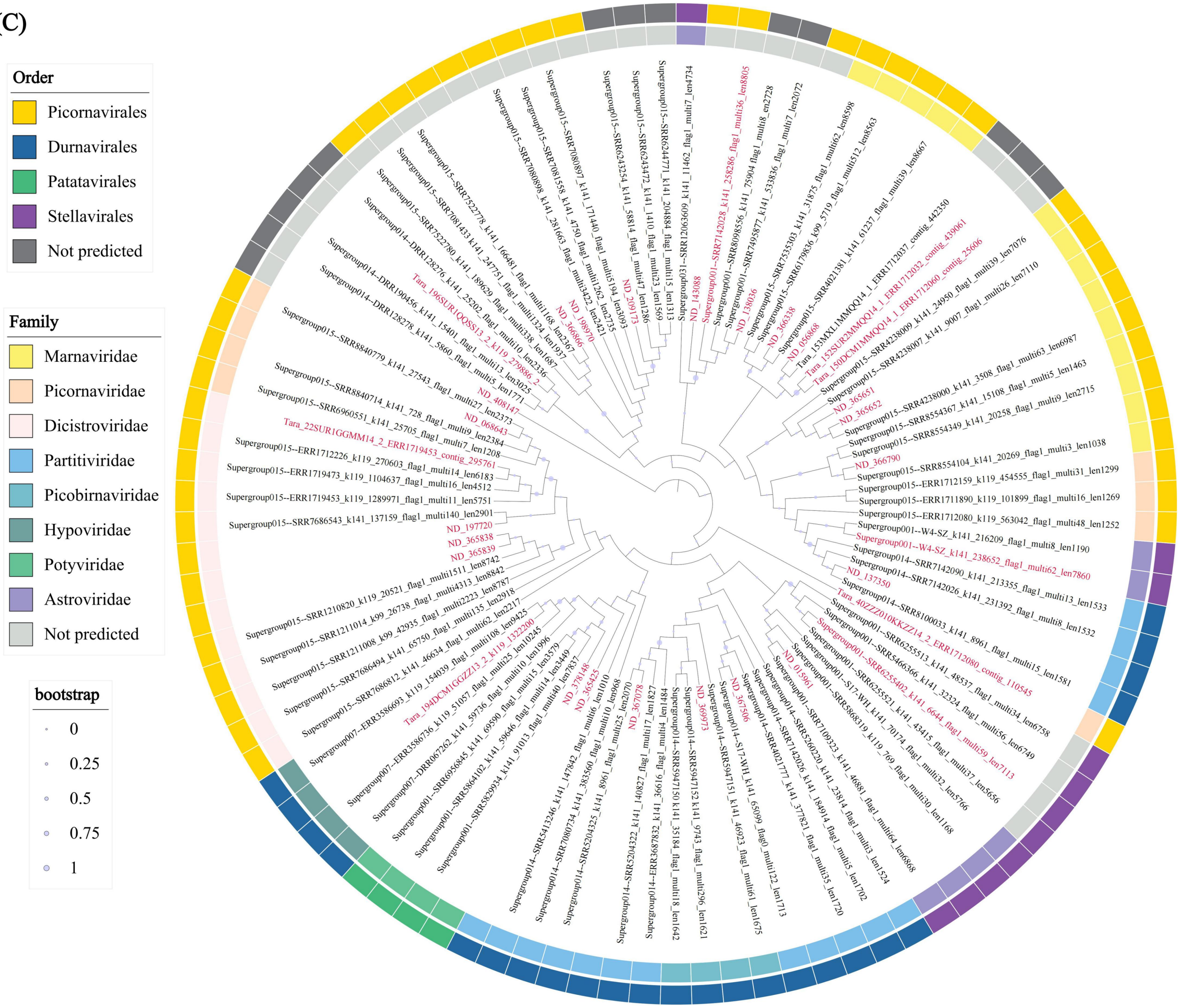


**Fig. S1 | Continue. (C)** Phylogenetic tree of RdRps for the Pisuviricota phylum. RNA viral RdRPs with AMGs are highlighted in red, while reference sequences are highlighted in black. The outer ring shows the order and family of RdRps, respectively.

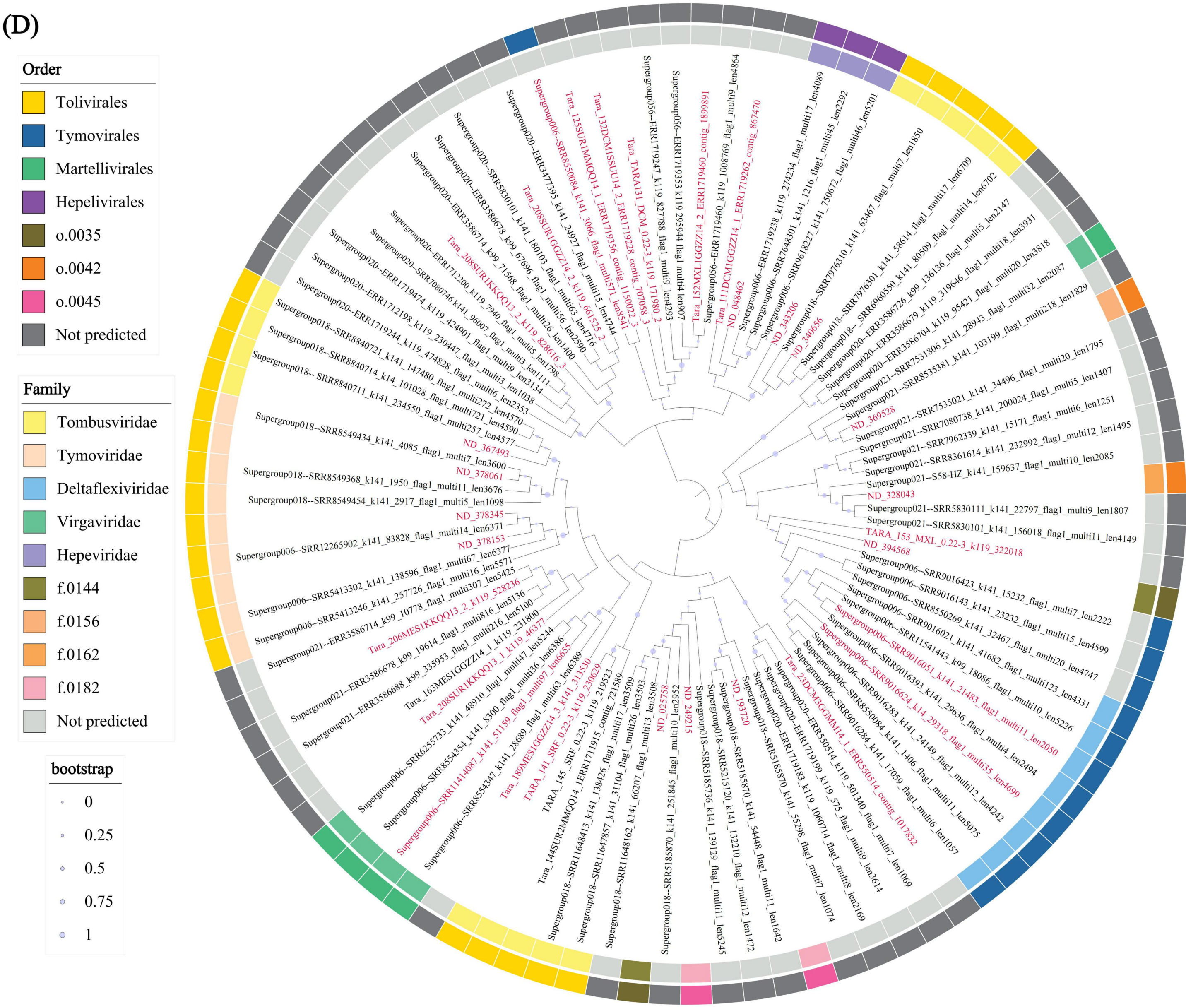


**Fig. S1 | Continue. (D)** Phylogenetic tree of RdRps for the Kitrinoviricota phylum. RNA viral RdRPs with AMGs are highlighted in red, while reference sequences are highlighted in black. The outer ring shows the order and family of RdRps, respectively.

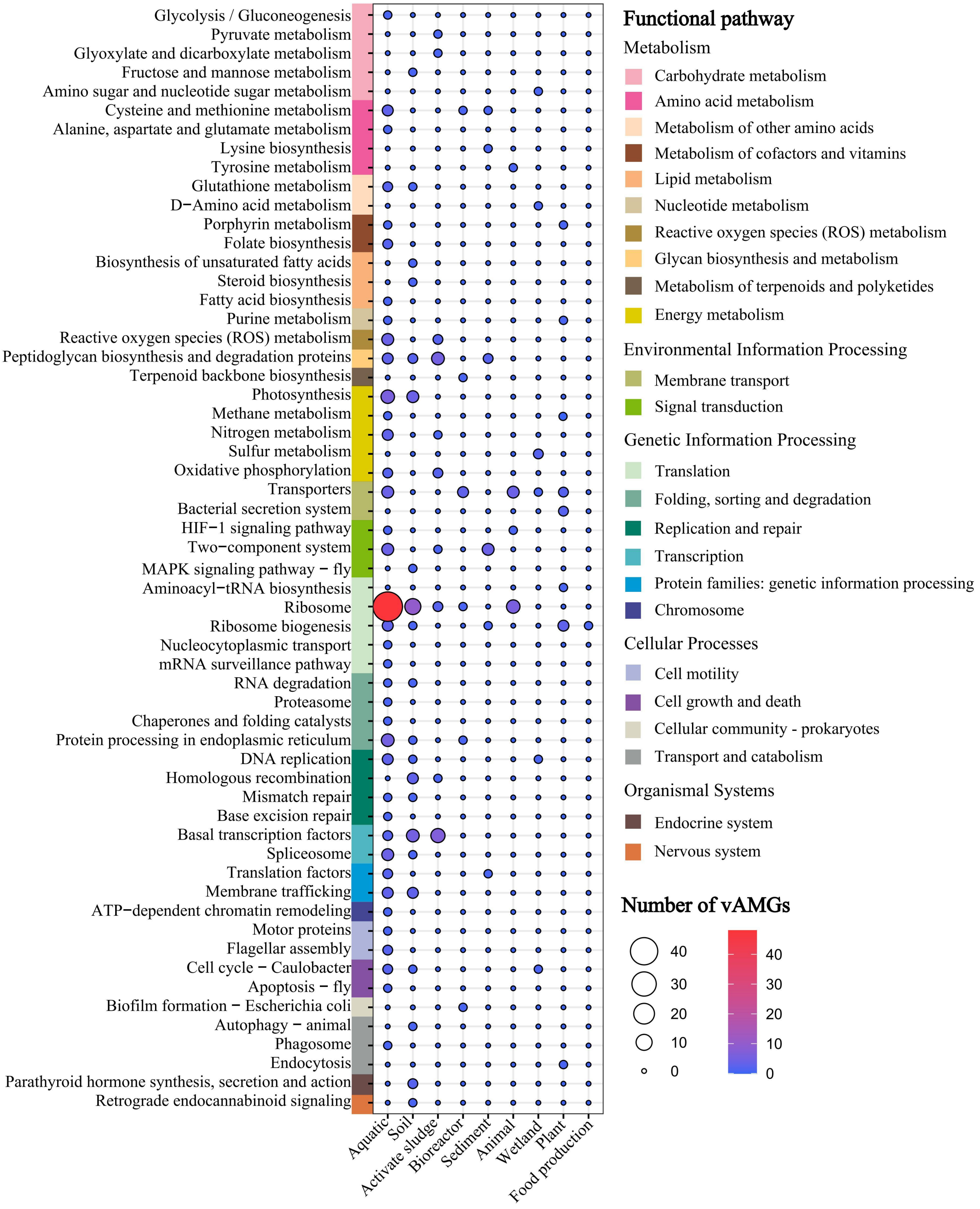


**Fig. S2 | RNA viral AMGs are distributed in nine habitats and the 58 biological pathways**.

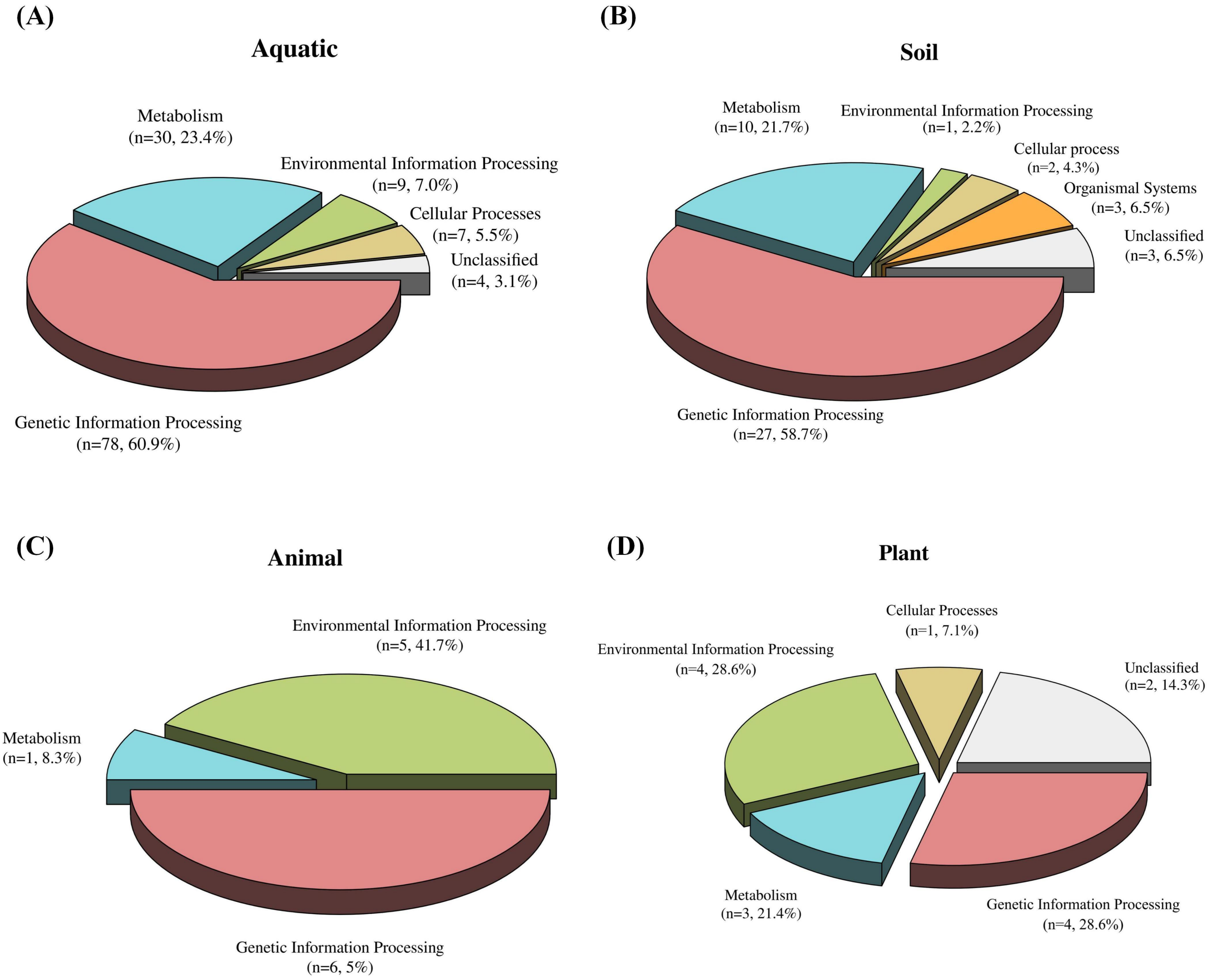


**Fig. S3 | Proportion of RNA vAMGs in each environment in metabolism, environmental information processing, genetic information processing, cellular processes, and biological systems.** The proportion of RNA vAMGs in aquatic **(A)**, soil **(B)**, animal **(C)**, plant **(D)** environments in metabolism, environmental information processing, genetic information processing, cellular processes and organismal systems.

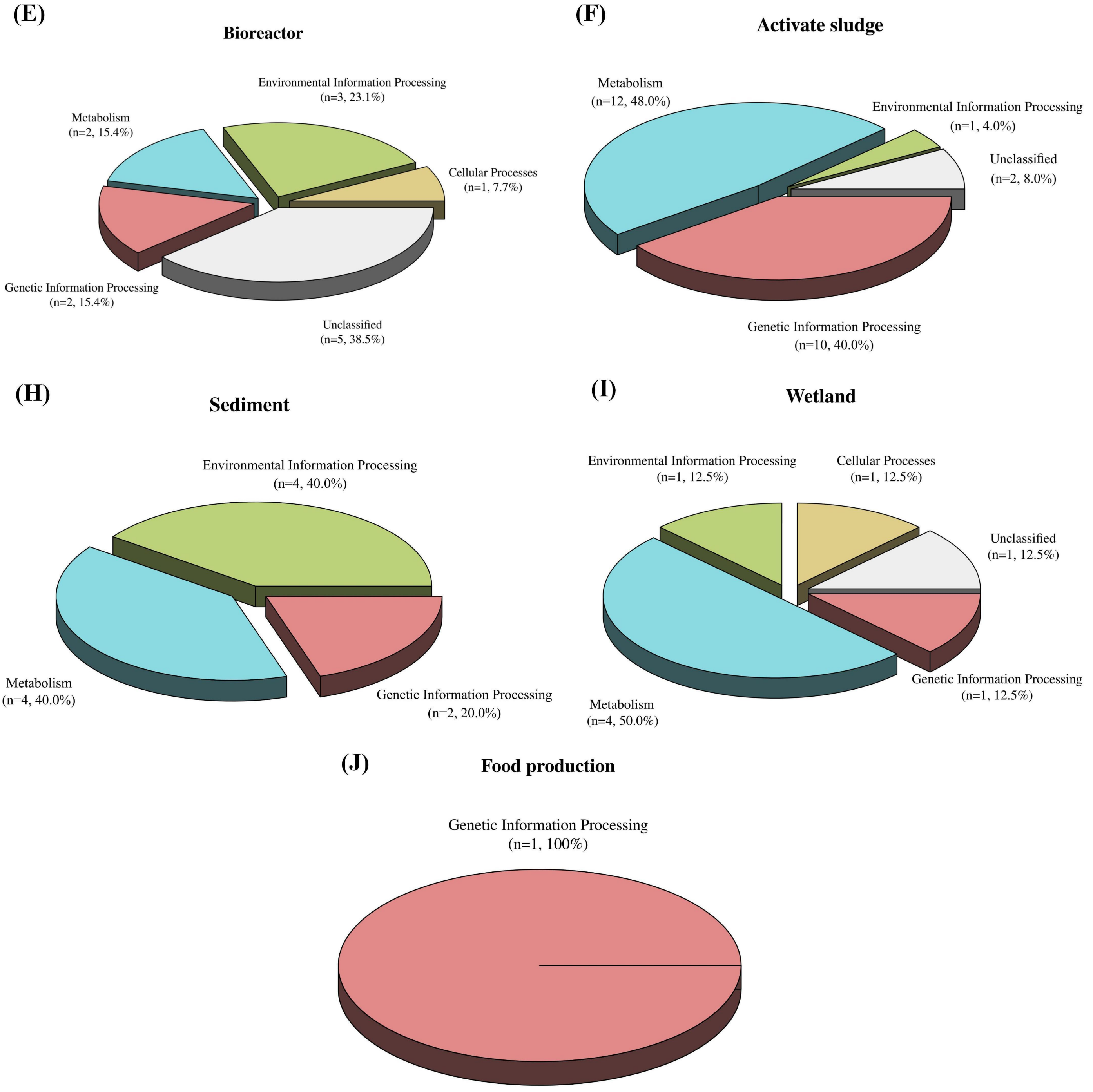


**Fig. S3 | Continue.** The proportion of RNA vAMGs in bioreactor **(E)**, activate sludge **(F)**, sediment **(H)**, wetland **(I)**, and food production **(J)** environments in metabolism, environmental information processing, genetic information processing, cellular processes and organismal systems.

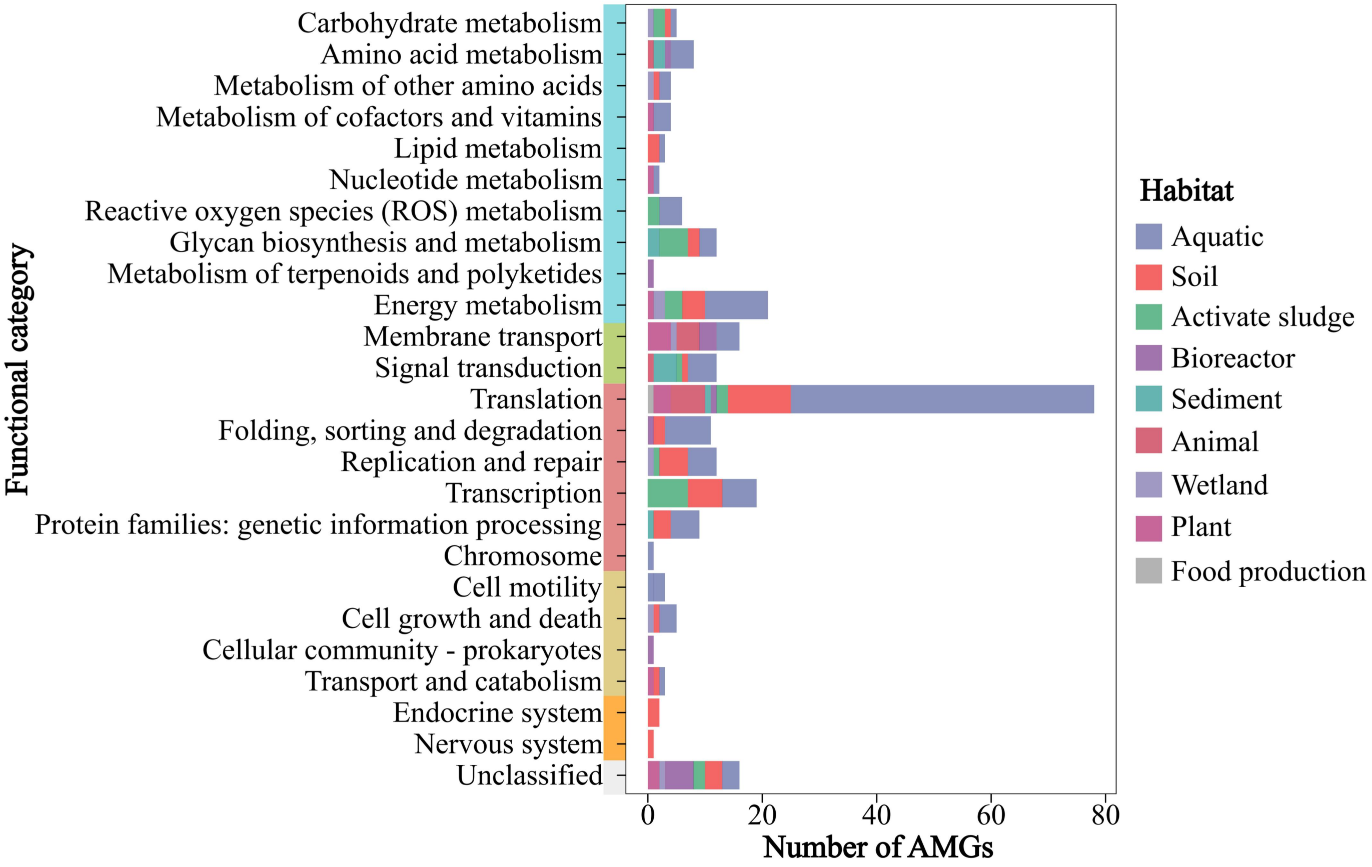


**Fig. S4 | Distribution of RNA vAMGs from different environments in 25 functional categories.** The y-axis colours from top to bottom represent metabolism, environmental information processing, genetic information processing, cellular processes, organismal systems and unclassified, consistent with the colour of Fig. S2.

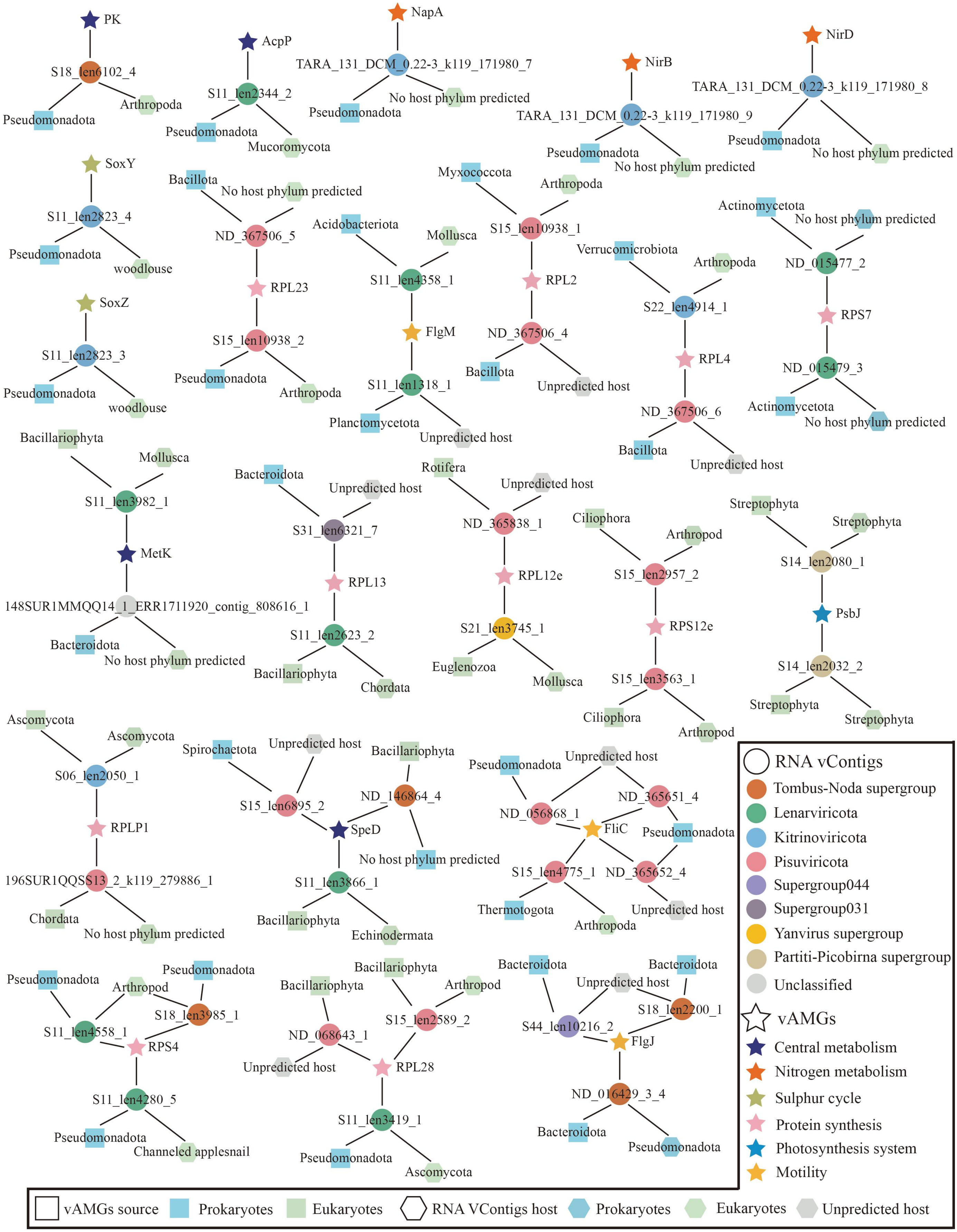


**Fig. S5 | Network diagram of RNA vContigs with host information and their AMGs sources, including vAMGs related to biogeochemical cycling, protein synthesis, and motility.** Circles represent RNA vContigs encoding AMGs, pentagons represent vAMGs, squares represent probable sources of vAMGs as predicted by the phylogenetic tree, and hexagons represent hosts that RNA vContigs may infect.

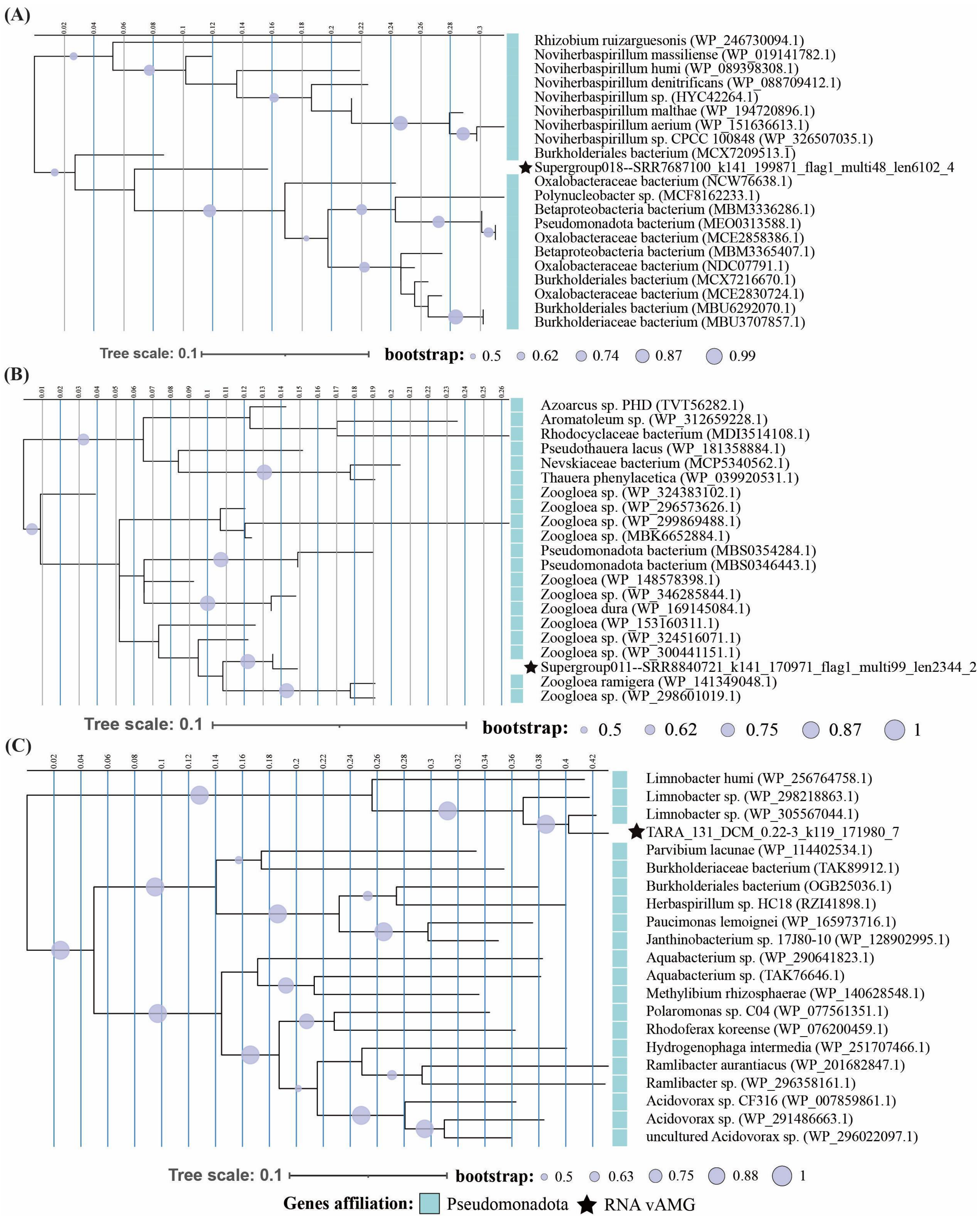


**Fig. S6 | Phylogenetic tree of RNA vAMGs that may have originated in prokaryotes.** **(A)** Phylogenetic tree of RNA viral *pk* and reference *pk* sequences found in NCBI nr database. **(B)** Phylogenetic tree of RNA viral *acpP* and reference *acpP* sequences found in NCBI nr database. **(C)** Phylogenetic tree of RNA viral *napA* and reference *napA* sequences found in NCBI nr database.

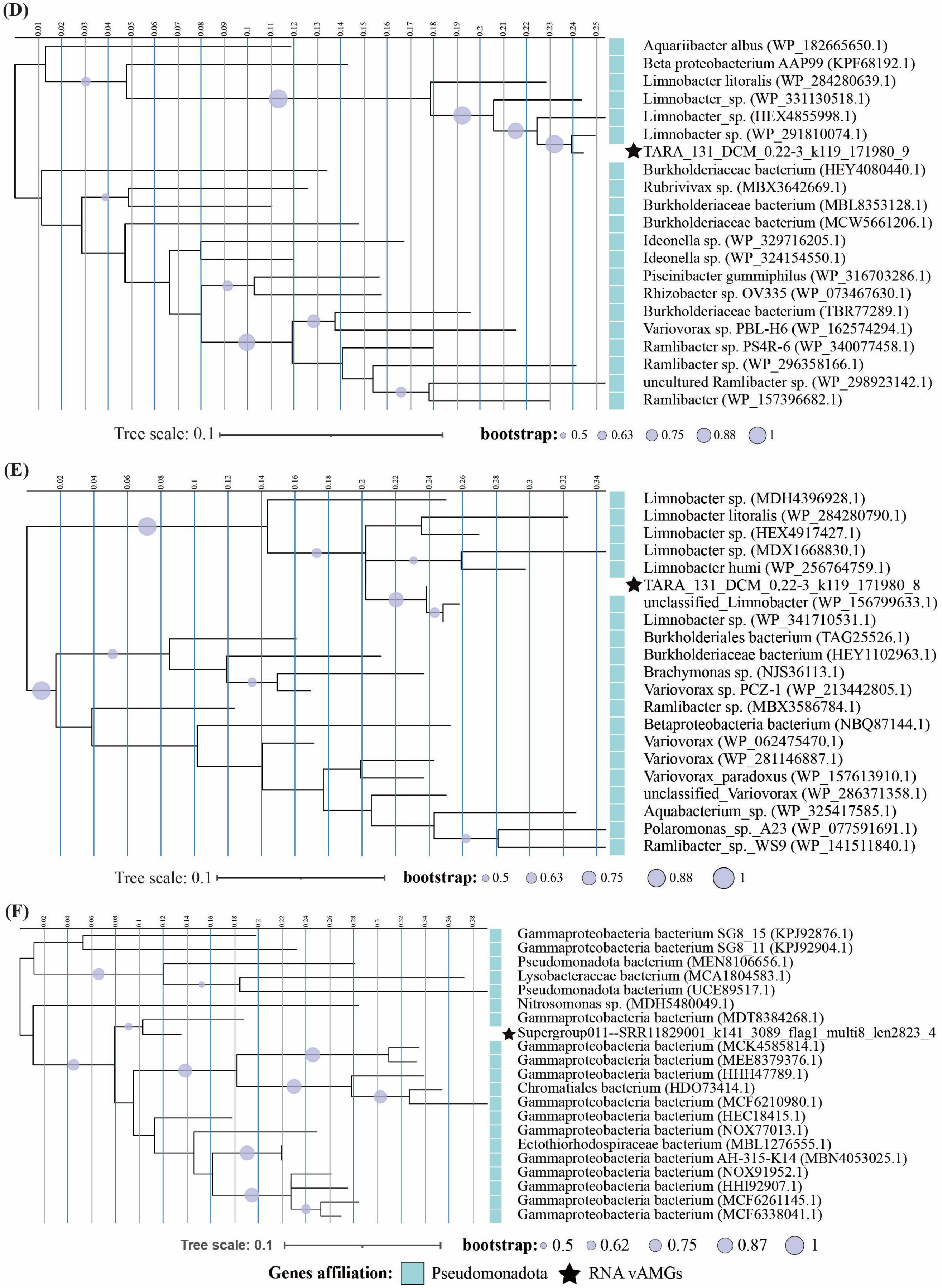


**Fig. S6 | Continue.** **(D)** Phylogenetic tree of RNA viral *nirB* and reference *nirB* sequences found in NCBI nr database. **(E)** Phylogenetic tree of RNA viral *nirD* and reference *nirD* sequences found in NCBI nr database. **(F)** Phylogenetic tree of RNA viral *soxY* and reference *soxY* sequences found in NCBI nr database.

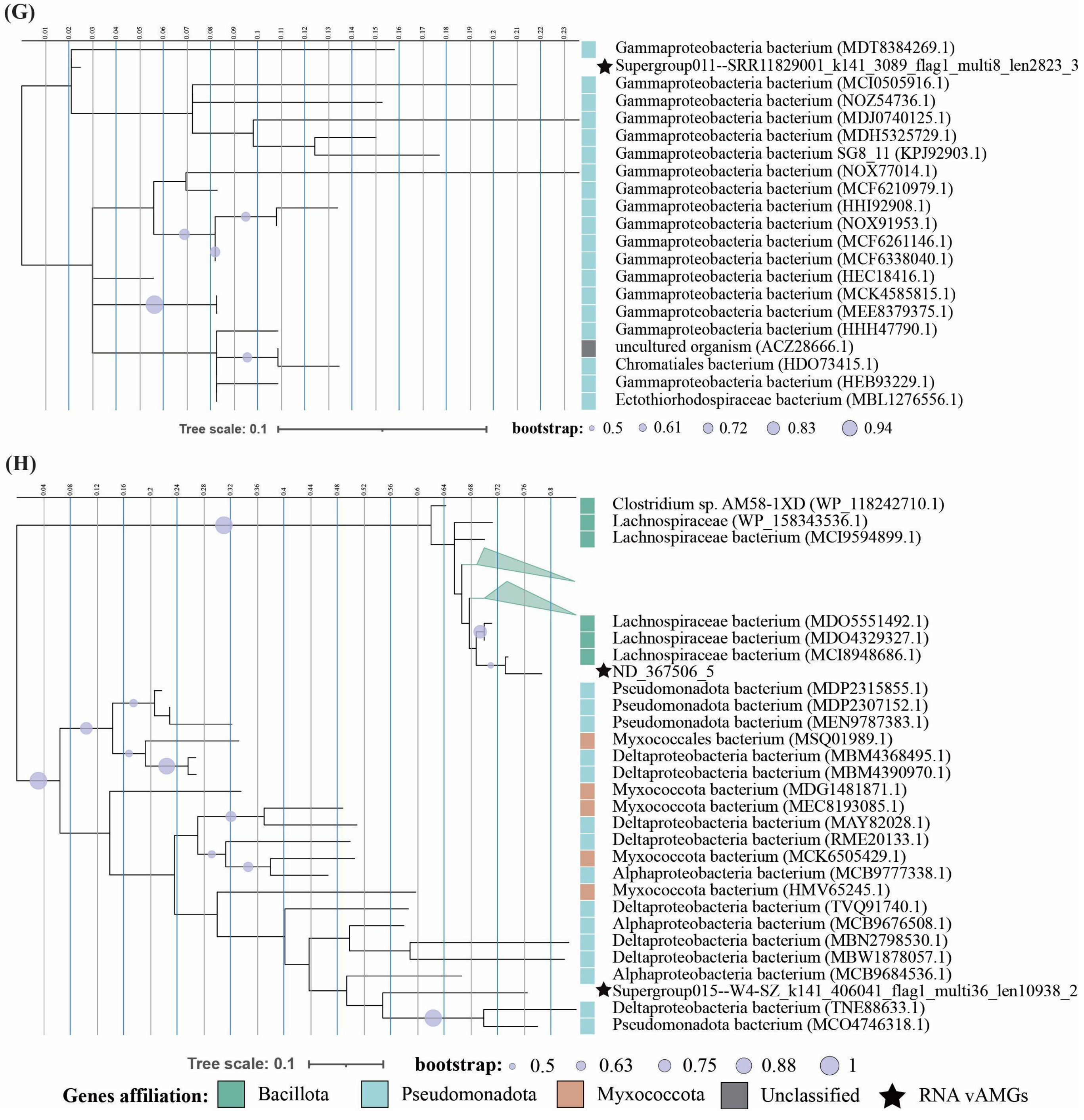


**Fig. S6 | Continue.** **(G)** Phylogenetic tree of RNA viral *soxZ* and reference *soxZ* sequences found in NCBI nr database. **(H)** Phylogenetic tree of RNA viral *rpl23* and reference *rpl23* sequences found in NCBI nr database.

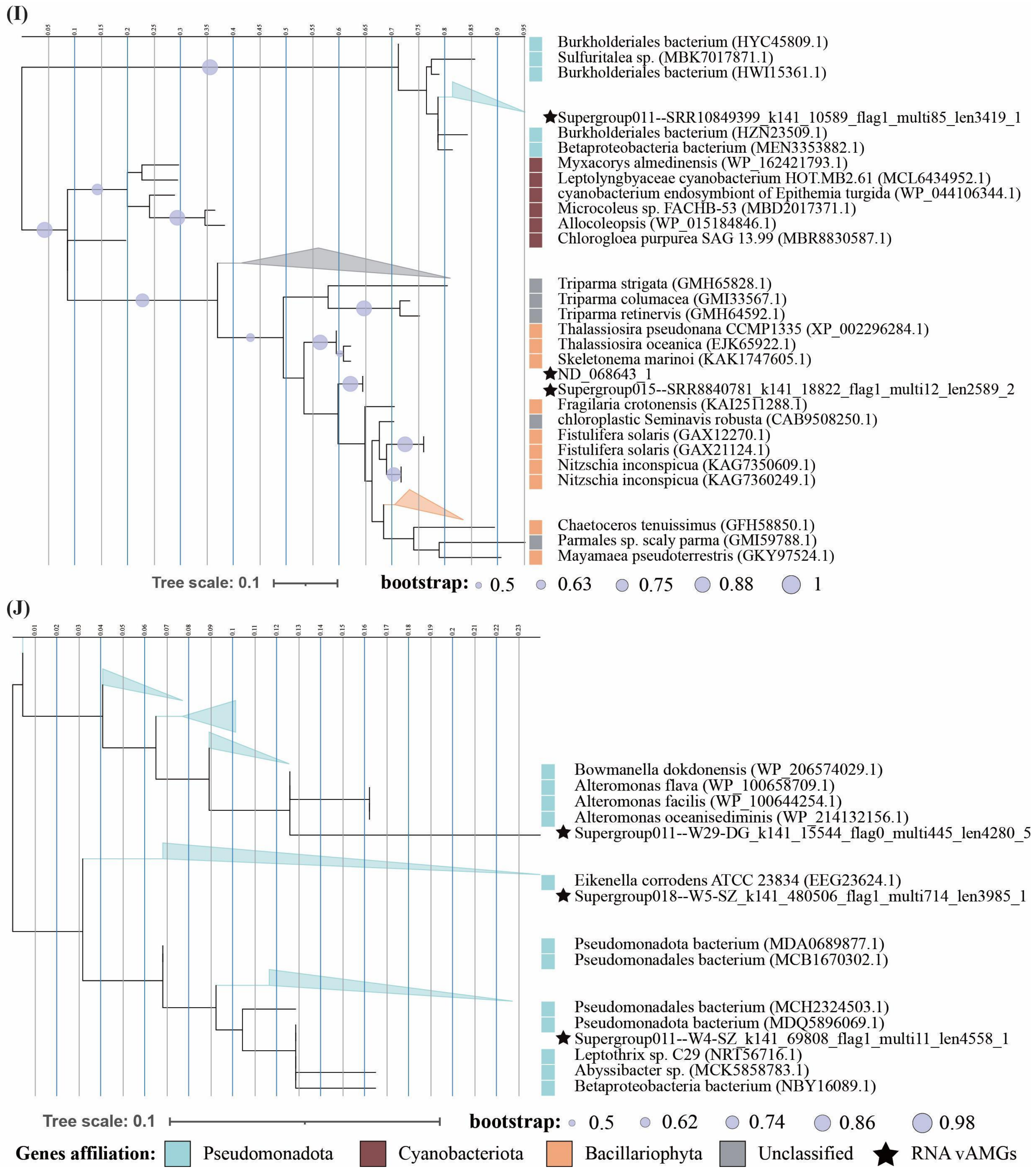


**Fig. S6 | Continue.** **(I)** Phylogenetic tree of RNA viral *rpl28* and reference *rpl28* sequences found in NCBI nr database. **(J)** Phylogenetic tree of RNA viral *rps4* and reference *rps4* sequences found in NCBI nr database.

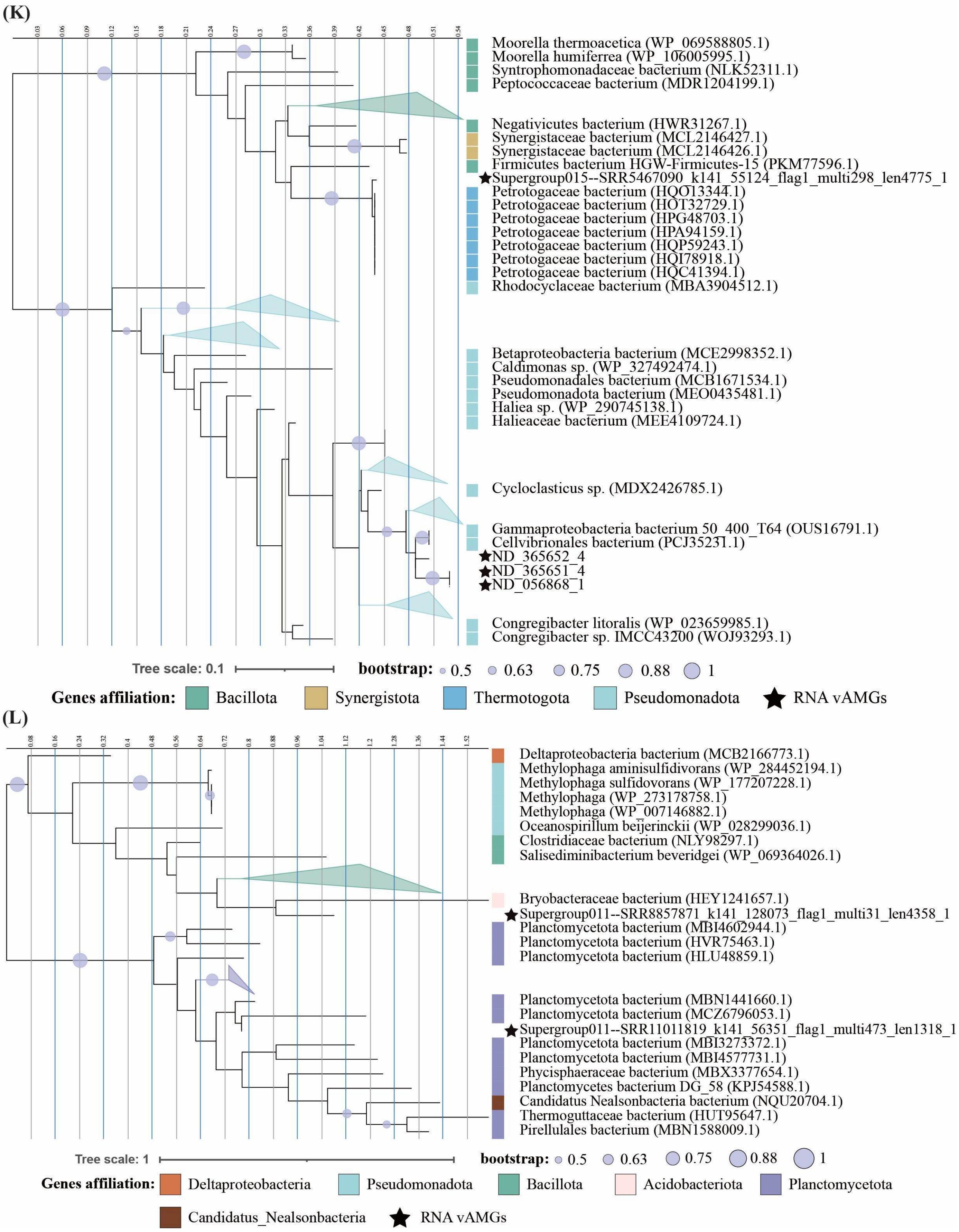


**Fig. S6 | Continue.** **(K)** Phylogenetic tree of RNA viral *fliC* and reference *fliC* sequences found in NCBI nr database. **(L)** Phylogenetic tree of RNA viral *flgM* and reference *flgM* sequences found in NCBI nr database.

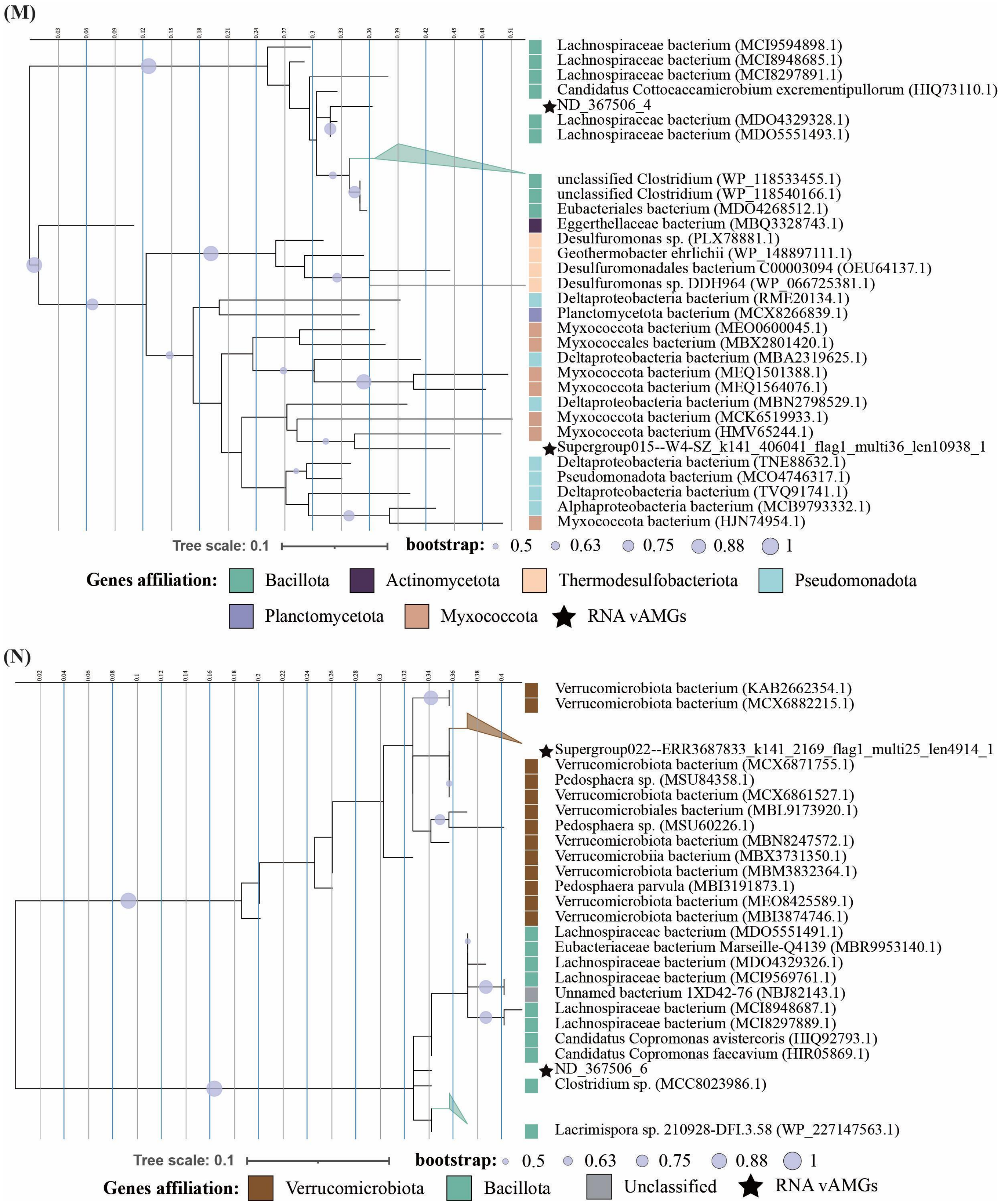


**Fig. S6 | Continue. (M)** Phylogenetic tree of RNA viral *rpl2* and reference *rpl2* sequences found in NCBI nr database. **(N)** Phylogenetic tree of RNA viral *rpl4* and reference *rpl4* sequences found in NCBI nr database.

**(O)**

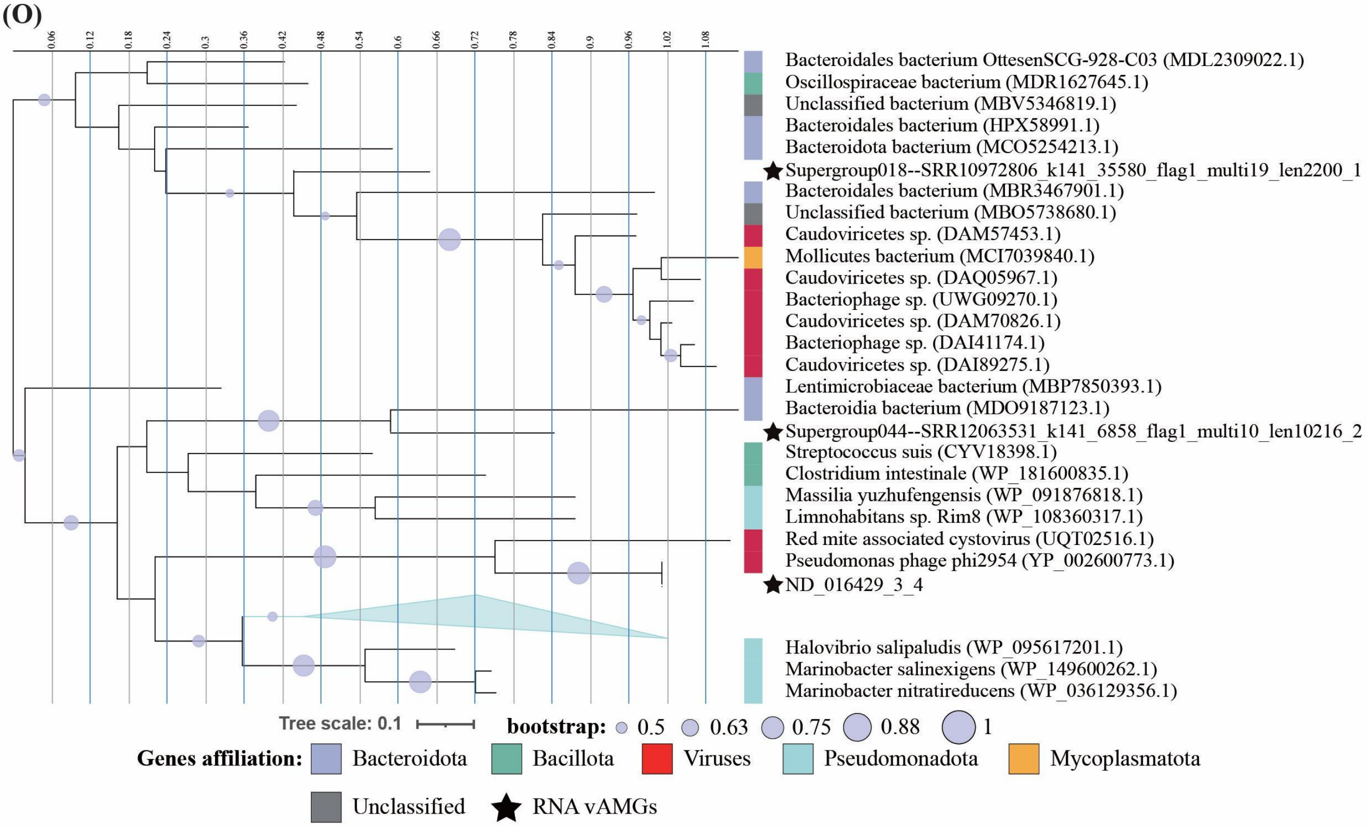


**(P)**

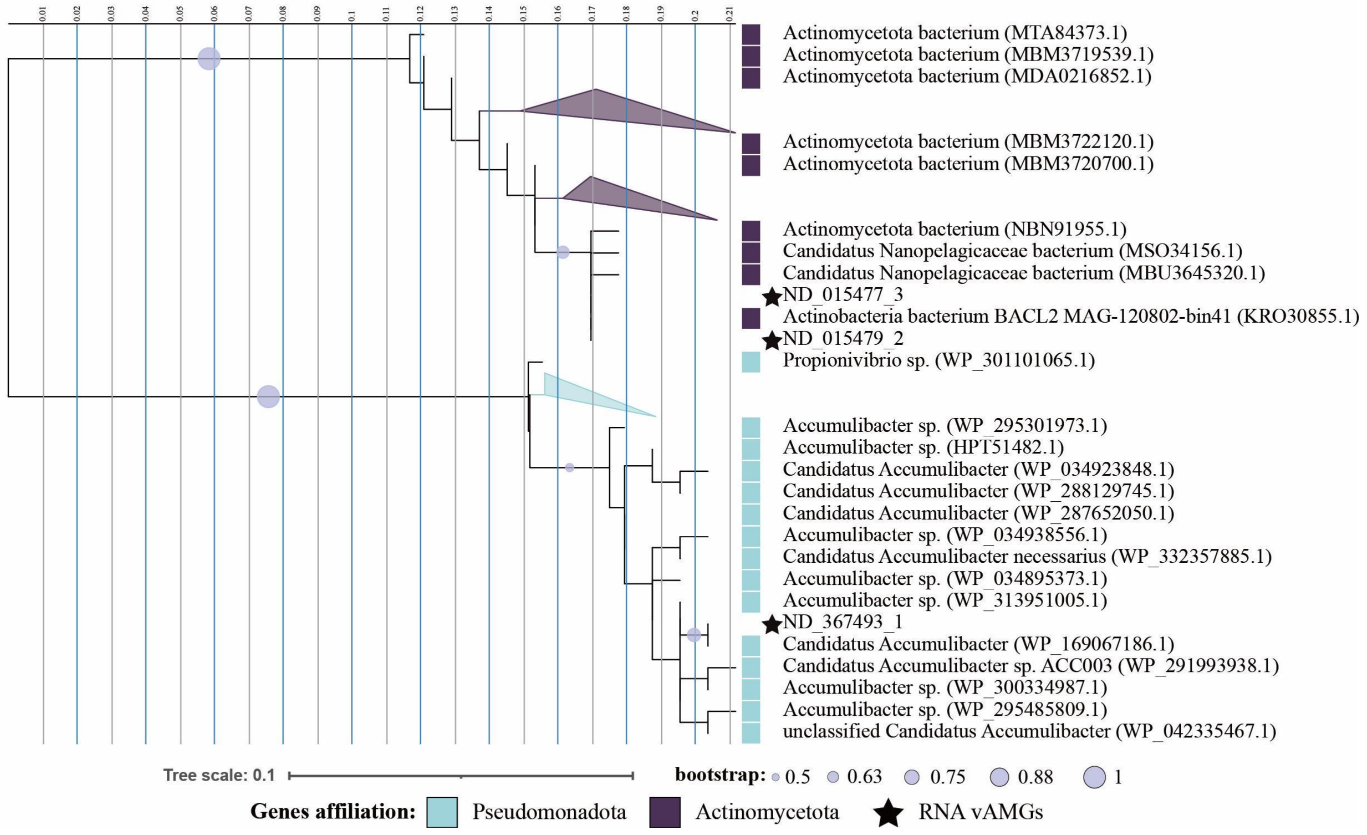


**Fig. S6 | Continue.** **(O)** Phylogenetic tree of RNA viral *flgJ* and reference *flgJ* sequences found in NCBI nr database. **(P)** Phylogenetic tree of RNA viral *rps12* and reference *rps12* sequences found in NCBI nr database.

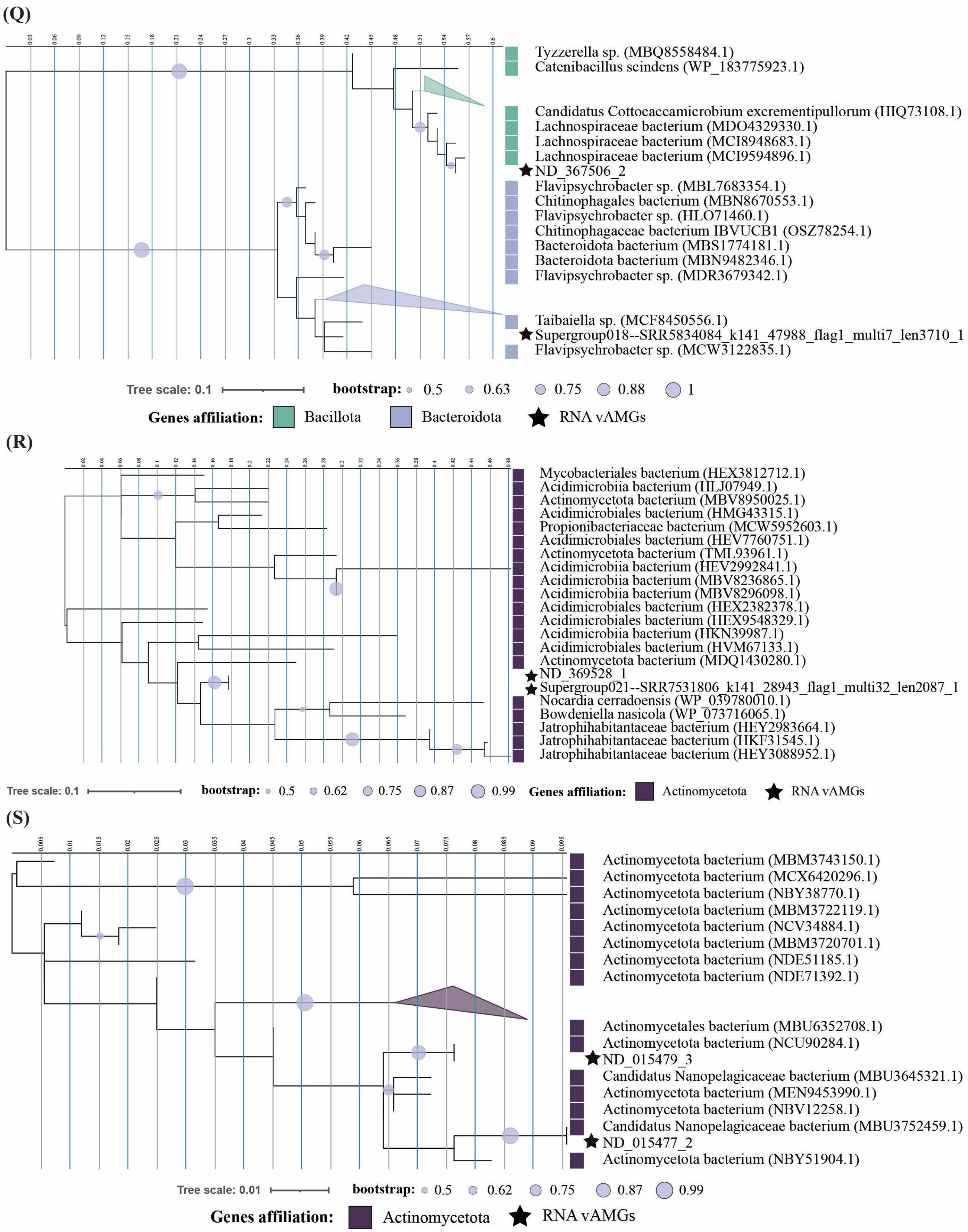


**Fig. S6 | Continue.** **(Q)** Phylogenetic tree of RNA viral *rpl22* and reference *rpl22* sequences found in NCBI nr database. **(R)** Phylogenetic tree of RNA viral *rpl36* and reference *rpl36* sequences found in NCBI nr database. **(S)** Phylogenetic tree of RNA viral *rps7* and reference *rps7* sequences found in NCBI nr database.

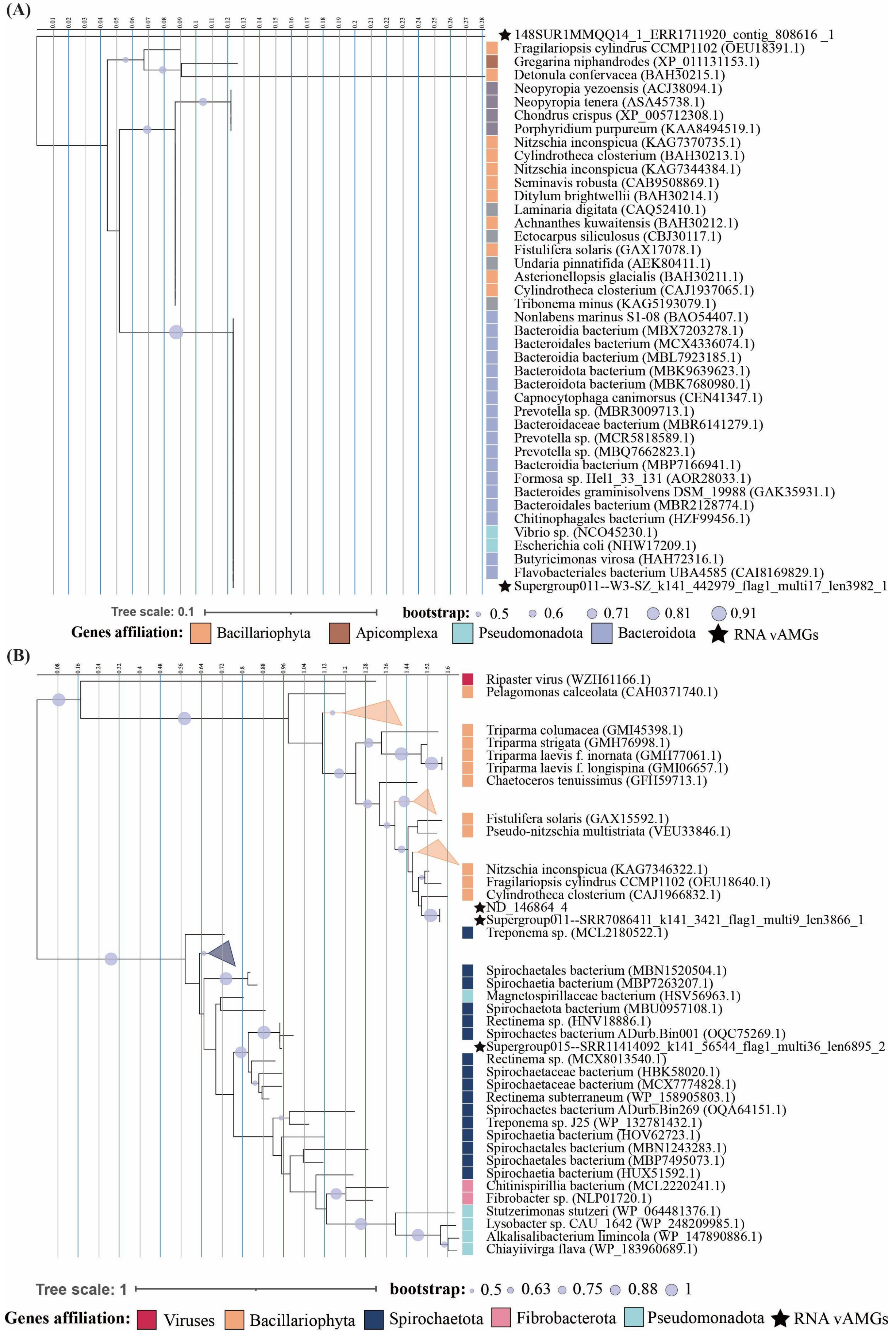


**Fig. S7 | Phylogenetic tree of RNA vAMGs that may have originated in eukaryotes.** **(A)** Phylogenetic tree of RNA viral *metK* and reference *metK* sequences found in NCBI nr database. **(B)** Phylogenetic tree of RNA viral *speD* and reference s*peD* sequences found in NCBI nr database.

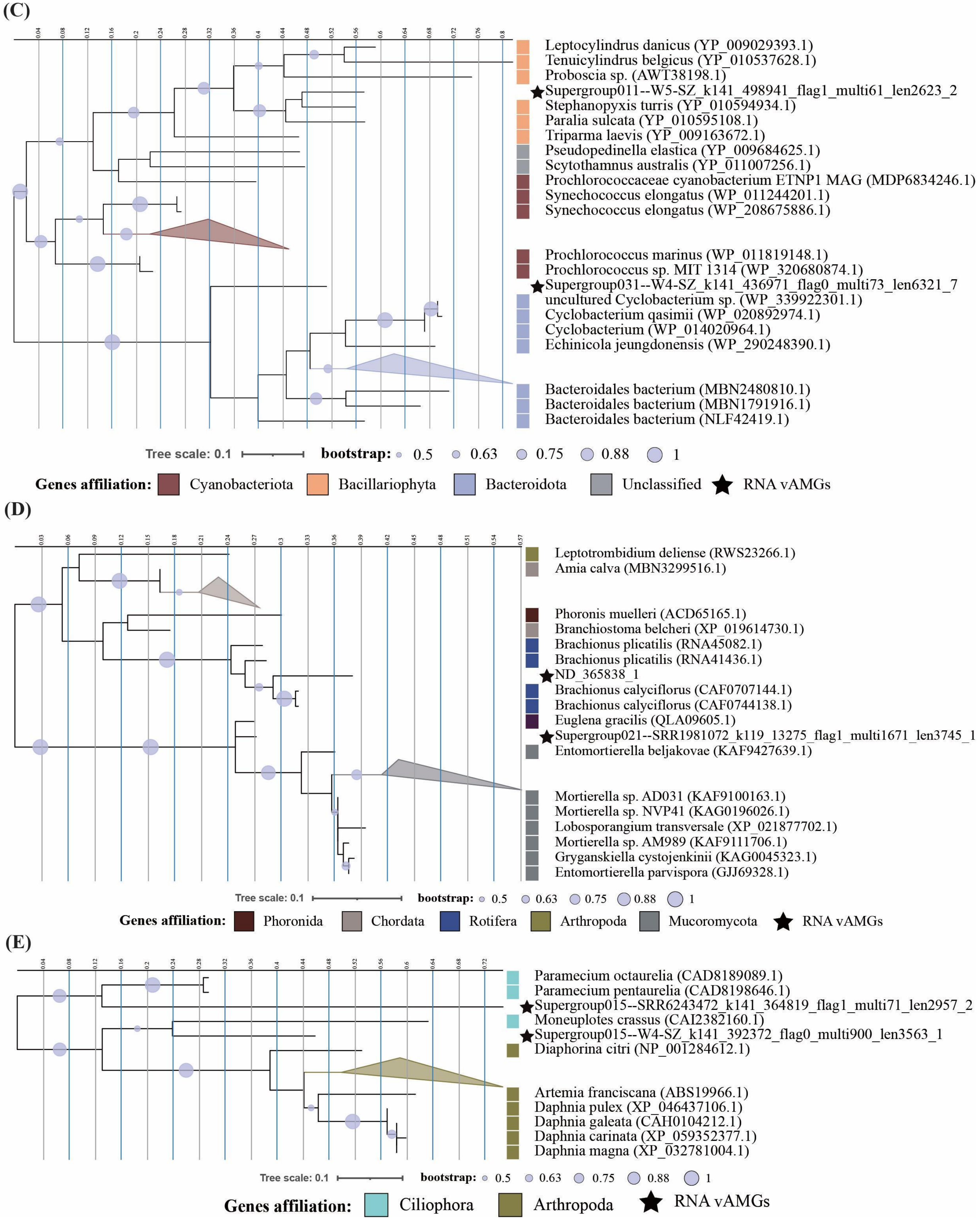


**Fig. S7 | Continue. (C)** Phylogenetic tree of RNA viral *rpl13* and reference *rpl13* sequences found in NCBI nr database. **(D)** Phylogenetic tree of RNA viral *rpl12e* and reference *rpl12e* sequences found in NCBI nr database. **(E)** Phylogenetic tree of RNA viral *rpS12e* and reference *rpS12e* sequences found in NCBI nr database.

(F)

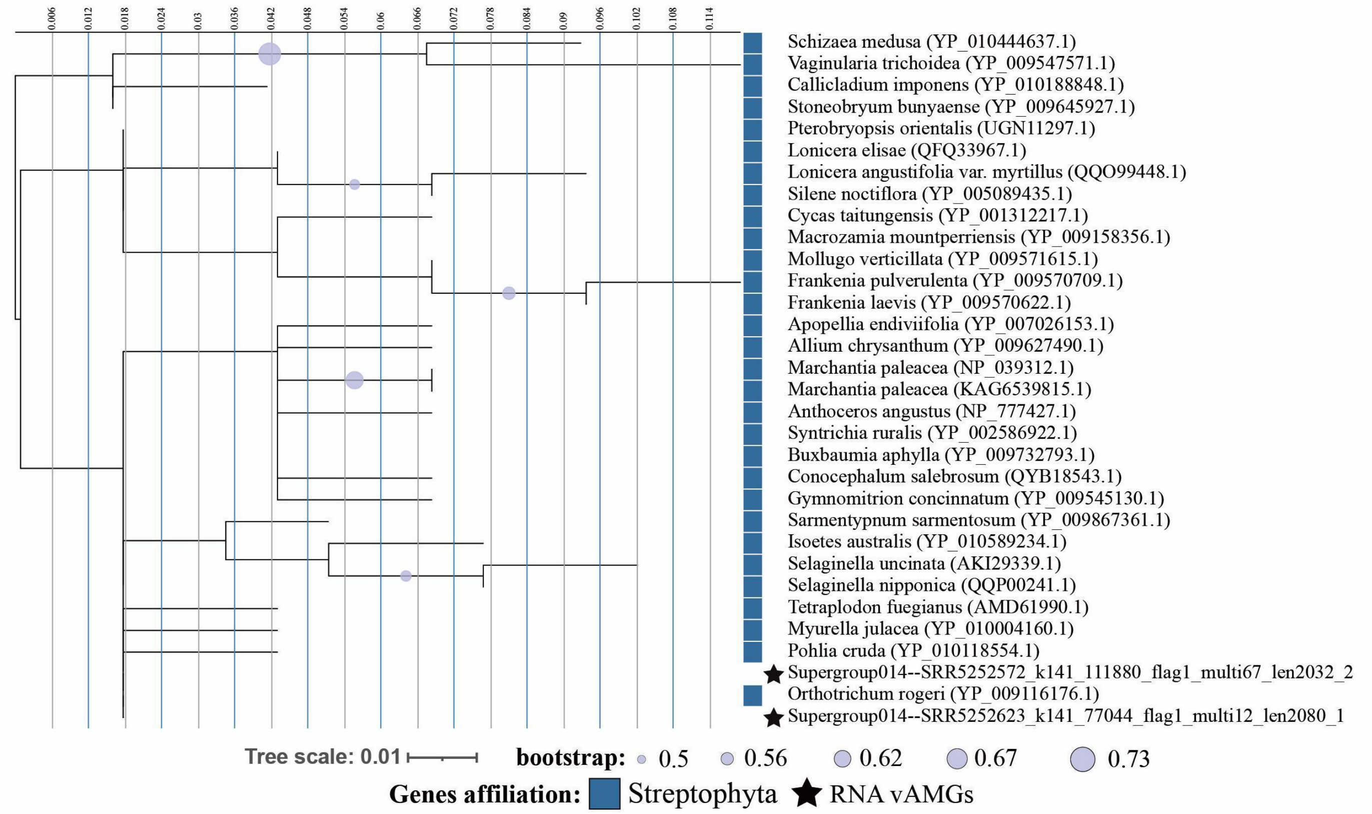


(J)

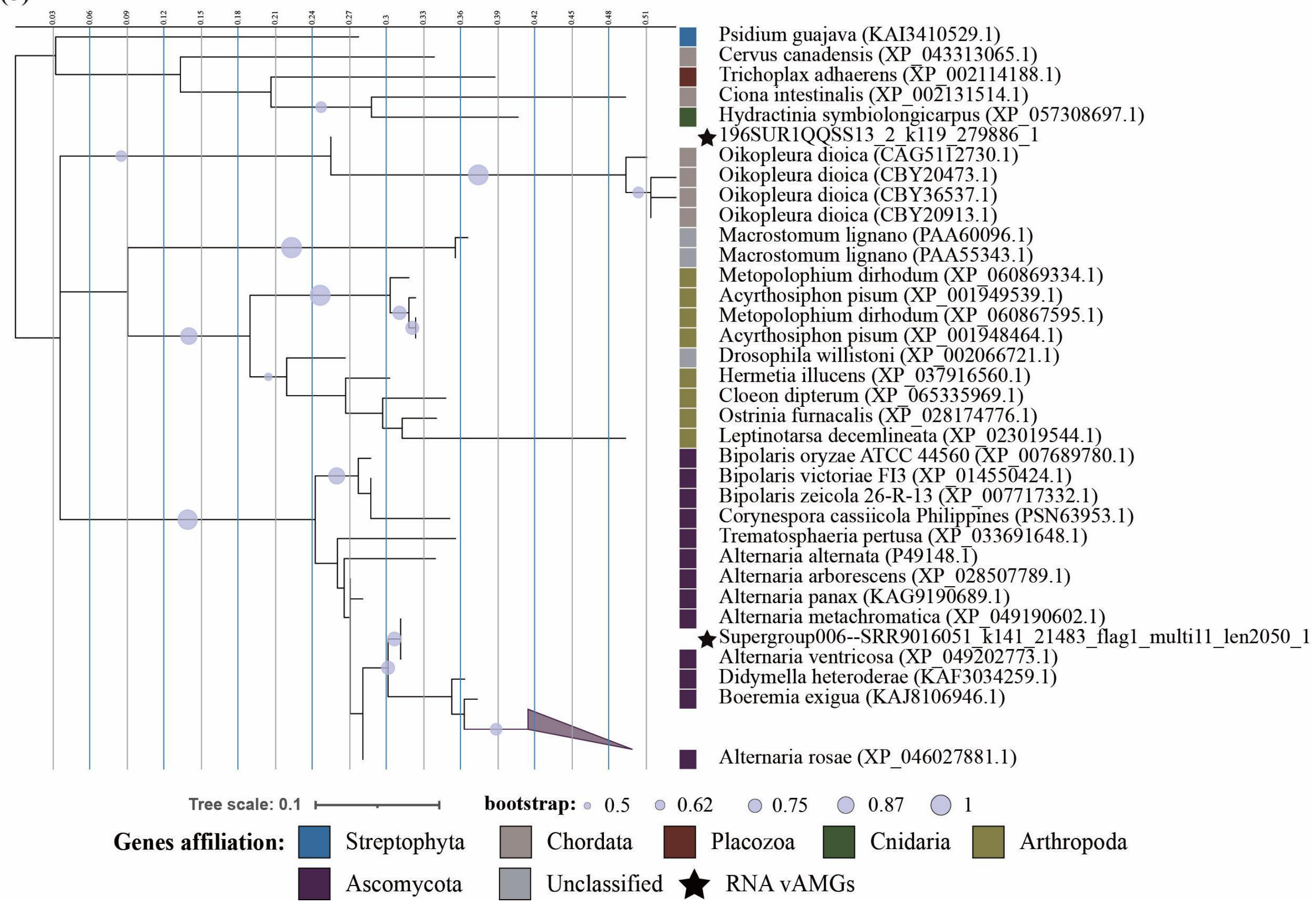


**Fig. S7 | Continue.** **(F)** Phylogenetic tree of RNA viral *psbJ* and reference *psbJ* sequences found in NCBI nr database. **(J)** Phylogenetic tree of RNA viral *rplP1* and reference *rplP1* sequences found in NCBI nr database.

(H)

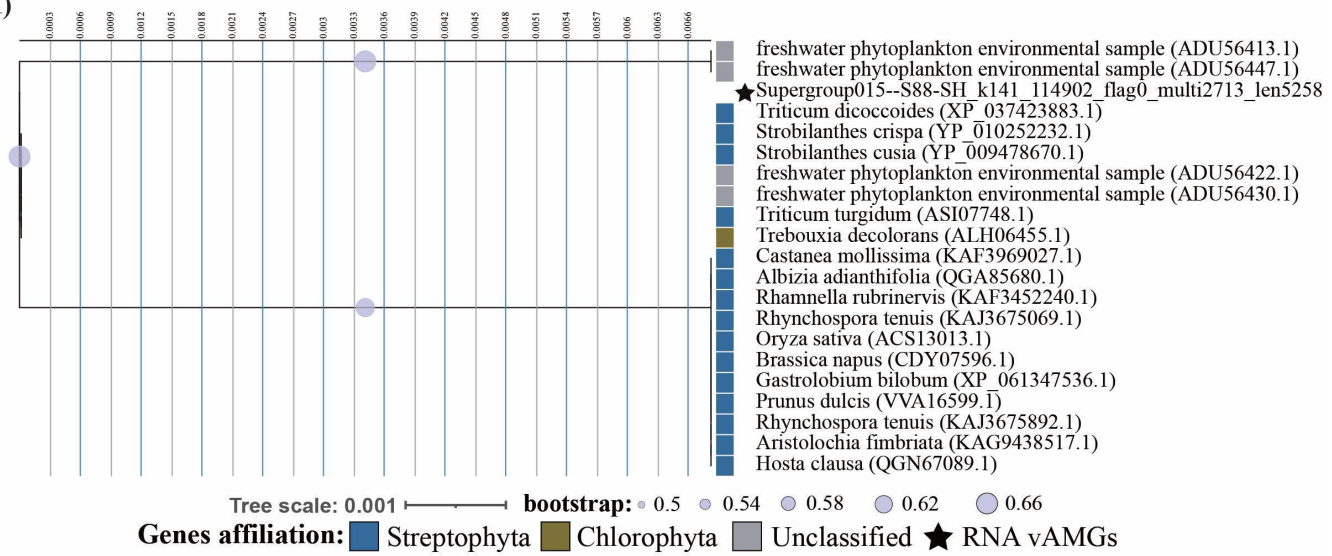


(I)

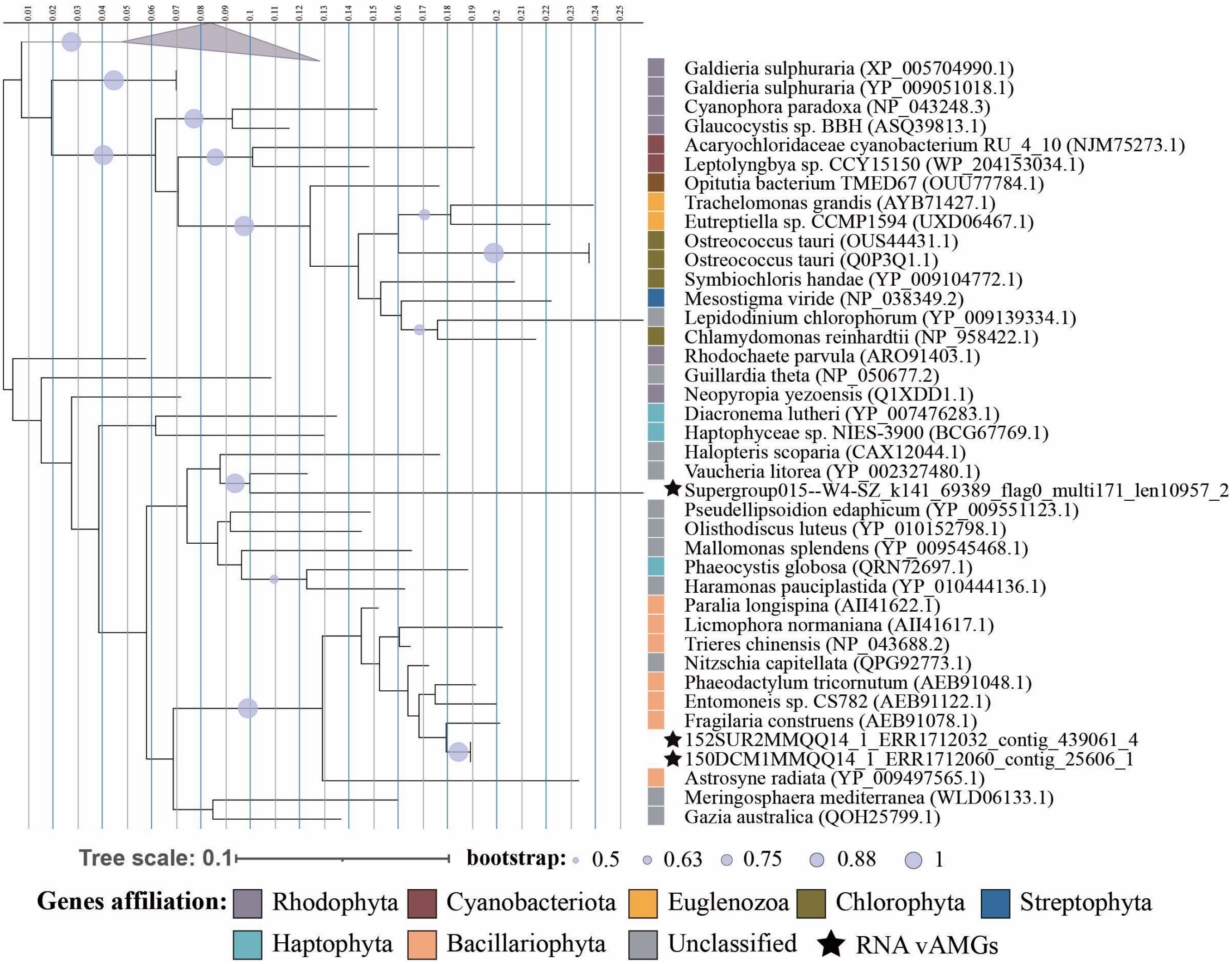


**Fig. S7 | Continue. (H)** Phylogenetic tree of RNA viral *psbA* and reference *psbA* sequences found in NCBI nr database. **(I)** Phylogenetic tree of RNA viral *psbC* and reference *psbC* sequences found in NCBI nr database.

(G)

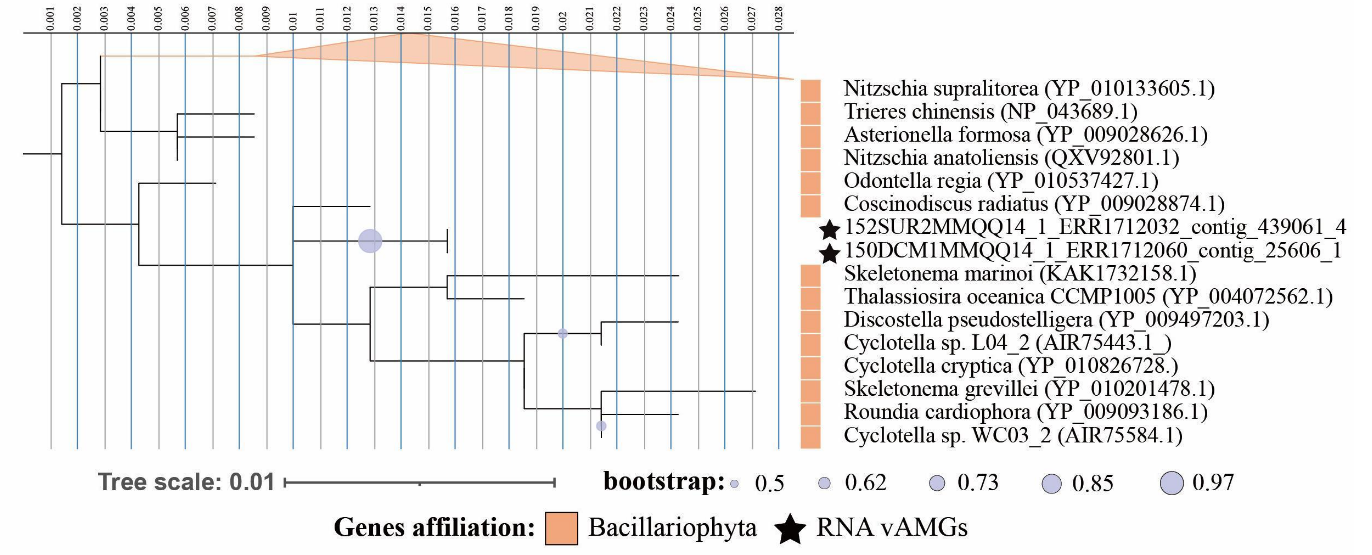


(K)

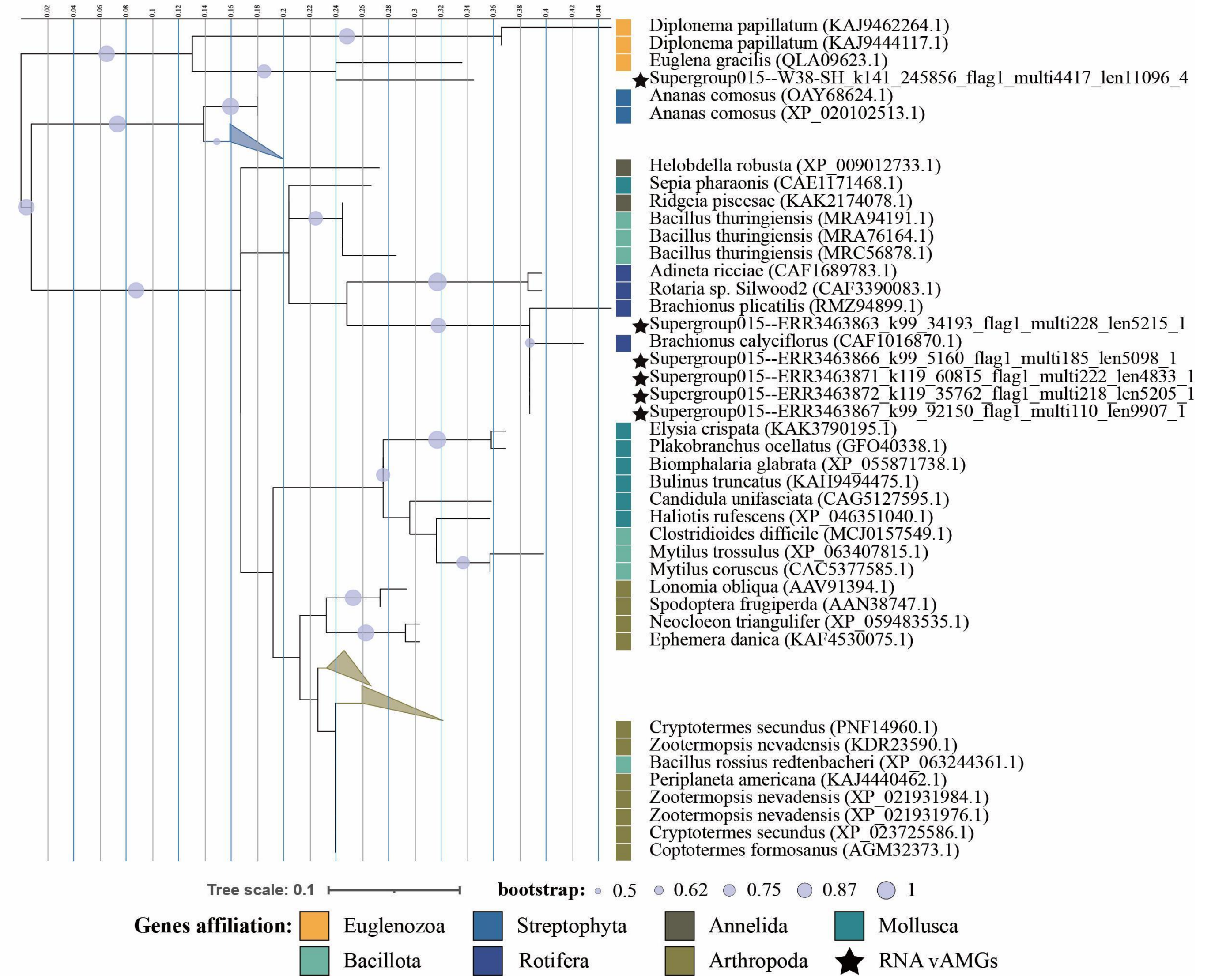


**Fig. S7 | Continue.** **(G)** Phylogenetic tree of RNA viral *psbD* and reference *psbD* sequences found in NCBI nr database. **(K)** Phylogenetic tree of RNA viral *rpl19e* and reference *rpl19e* sequences found in NCBI nr database.

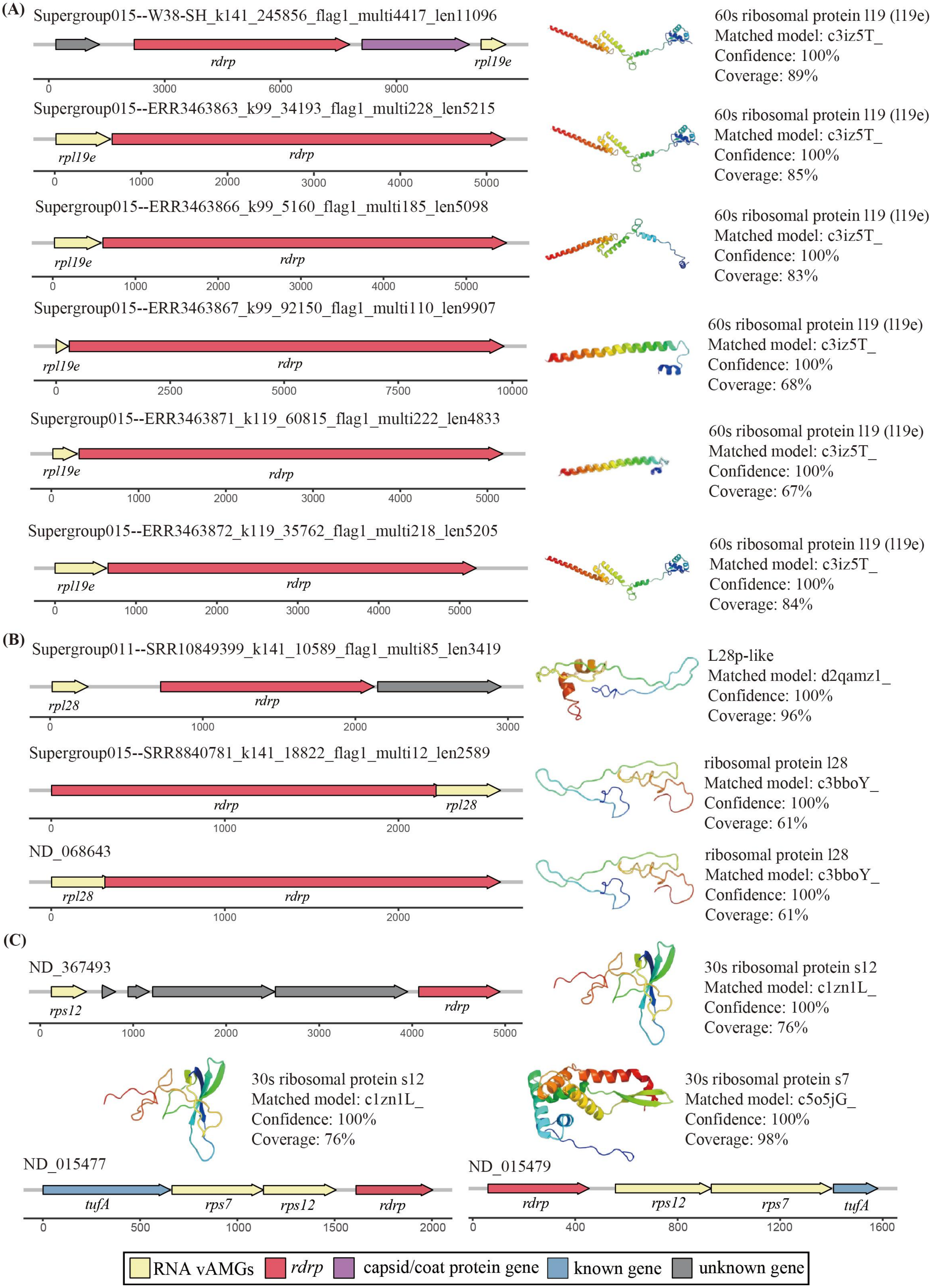


**Fig. S8 | Genome architecture of RNA vContig encoding a ribosomal protein gene and reference protein model.** **(A)** The genome architecture of RNA vContig encoding *rpl19e* gene and reference protein model for RPL19e. **(B)** Genome architecture of RNA vContig encoding *rpl28* gene and reference protein model for RPL28. **(C)** Genome architecture of RNA vContig encoding *rps12* gene and reference protein model for RPS12.

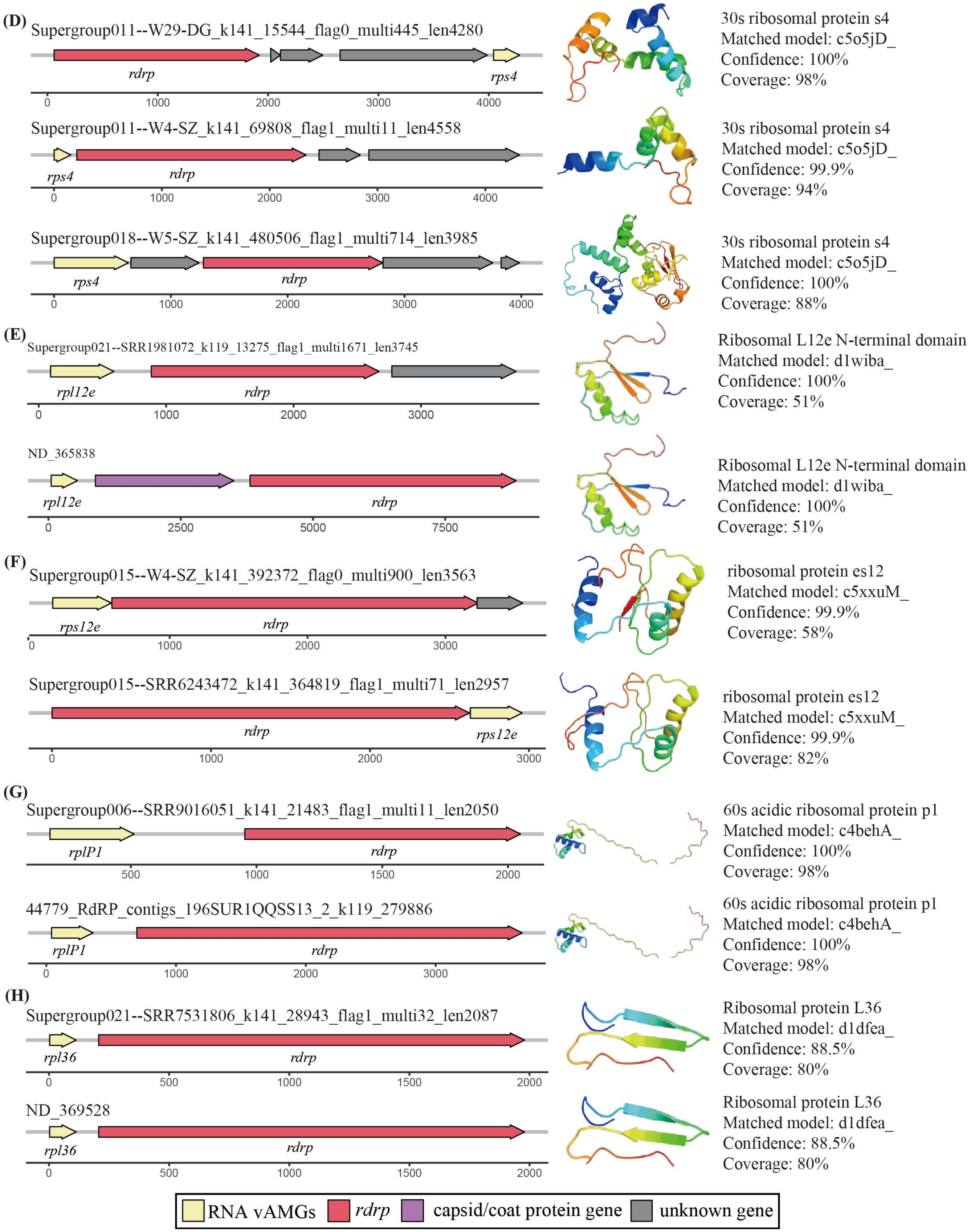


**Fig. S8 | Continue.** **(D)** Genome architecture of RNA vContig encoding *rps4* gene and reference protein model for RPS4. **(E)** Genome architecture of RNA vContig encoding *rpl28* gene and reference protein model for PRL28. **(F)** Genome architecture of RNA vContig encoding *rps12e* gene and reference protein model for RPS12e. **(G)** Genome architecture of RNA vContig encoding *rplP1* gene and reference protein model for LPLP1. **(H)** Genome architecture of RNA vContig encoding *rpl36* gene and reference protein model for RPL36.

(I)

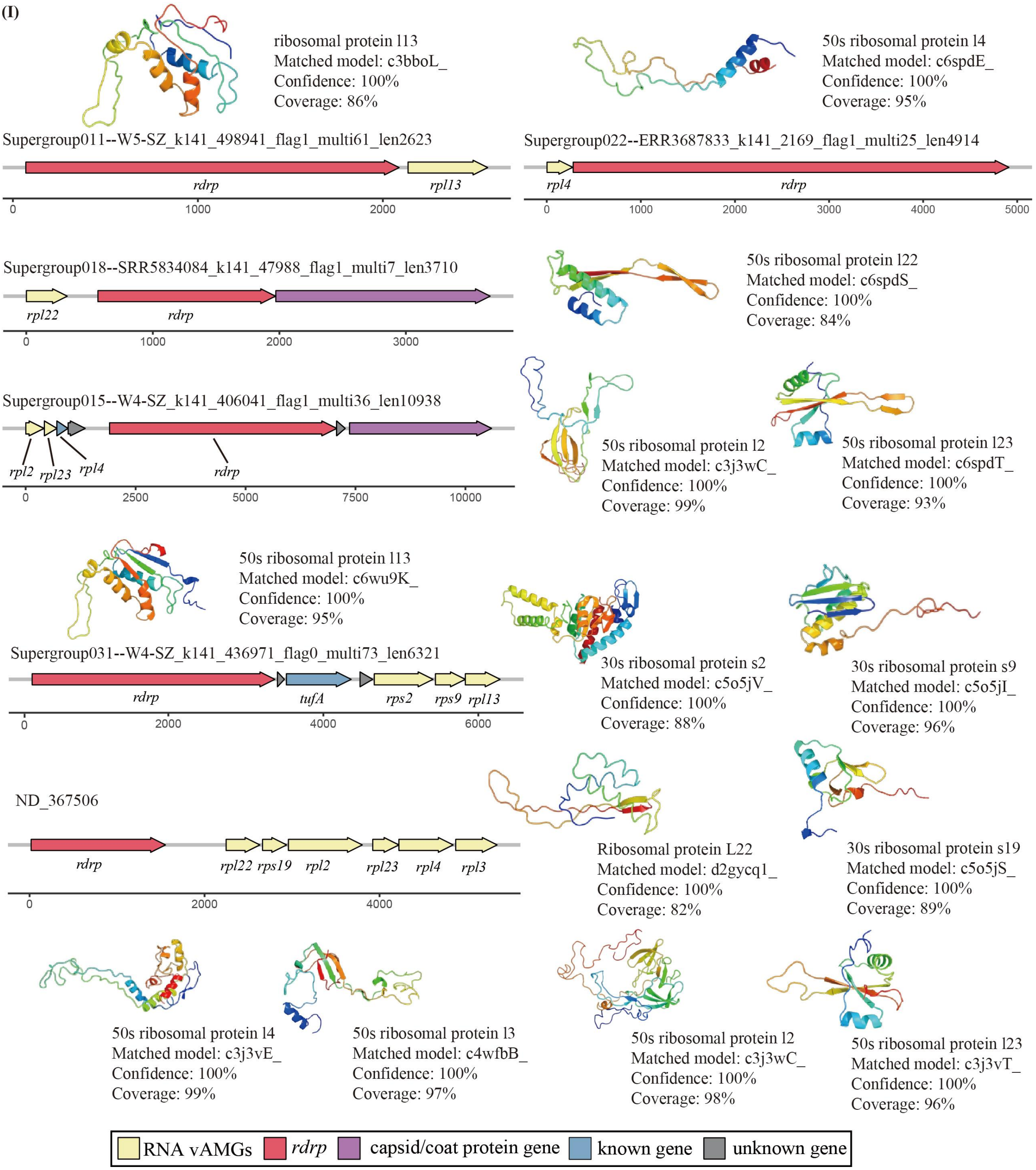


**Fig. S8 | Continue. (I)** Genome architecture of RNA vContig encoding *rpl13, rpl4, rpl22, rpl2, rpl23, rps2, rps9, rps19* and *rpl3* gene and reference protein model for RPL13, RPL4, RPL22, RPL2, RPL23, RPS2, RPS9, RPS19 and RPL3.
